# Deep phenotyping of the prostate tumor microenvironment reveals molecular stratifiers of relapse linked to inflammatory chemokine expression and aberrant metabolism

**DOI:** 10.1101/2024.05.13.593822

**Authors:** Sebastian Krossa, Maria K. Andersen, Elise Midtbust, Maximilian Wess, Antti Kiviaho, Abhibhav Sharma, Trond Viset, Øystein Størkersen, Guro F. Giskeødegård, Matti Nykter, Alfonso Urbanucci, Morten B. Rye, May-Britt Tessem

**Affiliations:** Department of Circulation and Medical Imaging, Norwegian University of Science and Technology; Central staff, St. Olavs Hospital HF, 7006, Trondheim, Norway; Clinic of Surgery, St. Olavs Hospital, Trondheim University Hospital, Trondheim, Norway; Prostate Cancer Research Center, Faculty of Medicine and Health Technology, Tampere University and TAYS Cancer Center, Tampere, Finland; HUNT Center for Molecular and Clinical Epidemiology, Department of Public Health and Nursing, Norwegian University of Science and Technology (NTNU), Trondheim, Norway; Department of Pathology, St. Olavs Hospital, Trondheim University Hospital, Trondheim, Norway; Foundation for the Finnish Cancer Institute, Helsinki, Finland; Department of Tumor Biology, Institute for Cancer Research, Oslo University Hospital, Oslo, Norway; Department of Clinical and Molecular Medicine, Norwegian University of Science and Technology

## Abstract

Understanding the molecular characteristics and changes of the tumor microenvironment (TME) associated with aggressive prostate cancer (PCa) is essential for precise diagnosis and treatment. We interrogated spatially resolved integrated transcriptomics and metabolomics data to build molecular strafiers discriminating patients with aggressive, potentially relapsing, and metastasizing PCa. We report a relapse associated (RA) gene expression signature characterized by upregulated immune response related gene expression scoring high in cancer, stroma, and glandular tissue of relapsing patients. Further, we identified a signature specific to a distinct sub-group of morphologically non-cancerous glands in prostate tissue from patients with relapsing cancer. This signature, named chemokine-enriched-gland (CEG) signature, was characterized by upregulated gene expression of pro-inflammatory chemokines. Glands with a high CEG score were enriched for club-like cells and surrounding stroma was infiltrated by immune cells. Tissue regions scoring high for both CEG and RA signatures were associated with reduced levels of citrate and zinc and loss of normal prostate secretory gland functions via reduced expression of genes necessary for citrate secretion. In summary we report that aggressive PCa is associated with an increased inflammatory status linked to chemokine production and club-like cell enrichment in potentially pre-cancerous prostate glands displaying an aberrant metabolism.

## Introduction/Main

Prostate cancer (PCa) is the most common malignancy among men in western countries [1]. This disease is clinically, morphologically, and molecularly highly heterogeneous, which determines aggressiveness and clinical outcome [2]. Approximately 30% of PCa patients experience relapse after prostatectomy and improving patient stratification based on clinical and molecular parameters is an ongoing effort [3]. In search for biomarkers that are predictive of relapse, substantial efforts are put into analyzing human PCa tissue having resulted in tissue-based biomarker assays such as Decipher and Prolaris [4]. More recently, to capture the transcriptomic heterogeneity single-cell and spatial transcriptomics (ST) have emerged as popular technologies to study PCa tissue samples [5-8]. ST has the advantage of capturing transcriptomics profiles in hundreds and thousands of locations across a tissue section thus offering a unique possibility to study the transcriptome in the spatial context, including intra-tumor heterogeneity, tumor microenvironment (TME) interactions, immune infiltration, and tissue-morphology-specific expression. Although single-cell transcriptomics data lacks spatial information, it has given new insight on cell-type specific transcriptome signatures present in the prostate [9, 10]. Applying such derived detailed cell-type signatures to ST data shows considerable potential, as it has revealed highly detailed information on tumor-immune interaction in the prostate TME [6].

Inflammation is a risk factor in cancer including PCa [11, 12]. Club cells can trigger inflammatory responses through chemokine secretion and are secretory cells initially characterized in the lung [13, 14], but have recently been observed in the benign prostate in a single-cell study [9]. In the healthy prostate, Club cells are located in the urothelial and proximal prostate zone, which was corroborated by studies in mice in which club cells were present during early prostate development and were proposed to contribute to benign hyperplasia by acting as multipotent progenitor cells [9, 15]. Subsequent single-cell RNA-sequencing and ST studies observed club cell populations in the prostate peripheral zone in association with PCa [5, 6]. These observations were also confirmed by the presence of Club cell markers in inflamed tissue adjacent to prostate tumor tissue [16]. The enrichment of Club cells in combination with reduced androgen receptor signaling in clinical tissue specimens has been shown as an effect of treatment with 5ARI and androgen deprivation therapy [17-19]. Inflammation and cancer lead to characteristic metabolic rewiring and the most prominent feature for PCa is the pronounced reduction of citrate levels compared to healthy prostate [20-22]. We have previously demonstrated that the diverse content of stroma can mask differences of metabolite and RNA levels between normal and cancerous PCa tissue in bulk analysis [23]. Here we used spatially resolved methodology integrating several -omics layers to gain an elevated understanding of PCa.

We integrated and analyzed multi-omics data including spatial and bulk transcriptomics and metabolomics data connected to detailed histopathology each collected from prostate tissue samples. We dissected the cancer biology of the TME from patients experiencing early post-radical prostatectomy relapse and from patients remaining relapse-free 10 years post-surgery. We found two gene expression signatures with elevated activity, club-like cell enrichment, and PCa-characteristic transcriptional and metabolic changes associated with these signatures in relapsing patients. Our approach allowed us to identify transcriptomic markers for highly localized metabolic aberrations in morphologically benign appearing glands in the TME of aggressive PCa. This highlights the substantial potential of integrating spatial-omics data to unravel the complexity and heterogeneity of cancer tissue.

## Results

To explore the gene expression in PCa tissue in its spatial, histology context we employed 10x Genomics’ spatial transcriptomics (ST) assay to collect data in 19854 circular spots from 32 tissue samples from prostatectomies of 8 PCa patients (**Figure 1a**). Of the 8 patients, 5 experienced relapse within 0 to 37 months and 3 remained relapse free during a follow-up of 10 years although they had similar clinicopathological features at diagnosis (**Table 1, Supplementary Table 1**). The 32 tissue samples were labeled according to sample location (cancer, field, or normal sample location, **Figure 1a**). Each ST spot was assigned a histopathology class, which included cancer with International Society for Urological Pathology (ISUP) grade group, non-cancerous glands (NCG), stroma, lymphocytes, lymphocyte enriched stroma and cancerous perineural invasion (PNI) obtained through hematoxylin-eosin(-saffron) (HE[S]) staining (**Figure 1a, Supplementary Figure 1a, Methods**). Each of the 32 samples was also subjected to multi-layered spatial analysis using immunohistochemistry (IHC) staining of bacterial antigens and mass spectrometry imaging (MSI) to obtain spatial metabolomics data. Further, these 32 samples together with an additional 142 samples from the remaining 29 of the 37 PCa patients were subjected to bulk transcriptomic and metabolomic analysis using RNA sequencing and high-resolution magic-angle spinning (HRMAS) NMR (**Figure 1a**).

**Figure 1.**
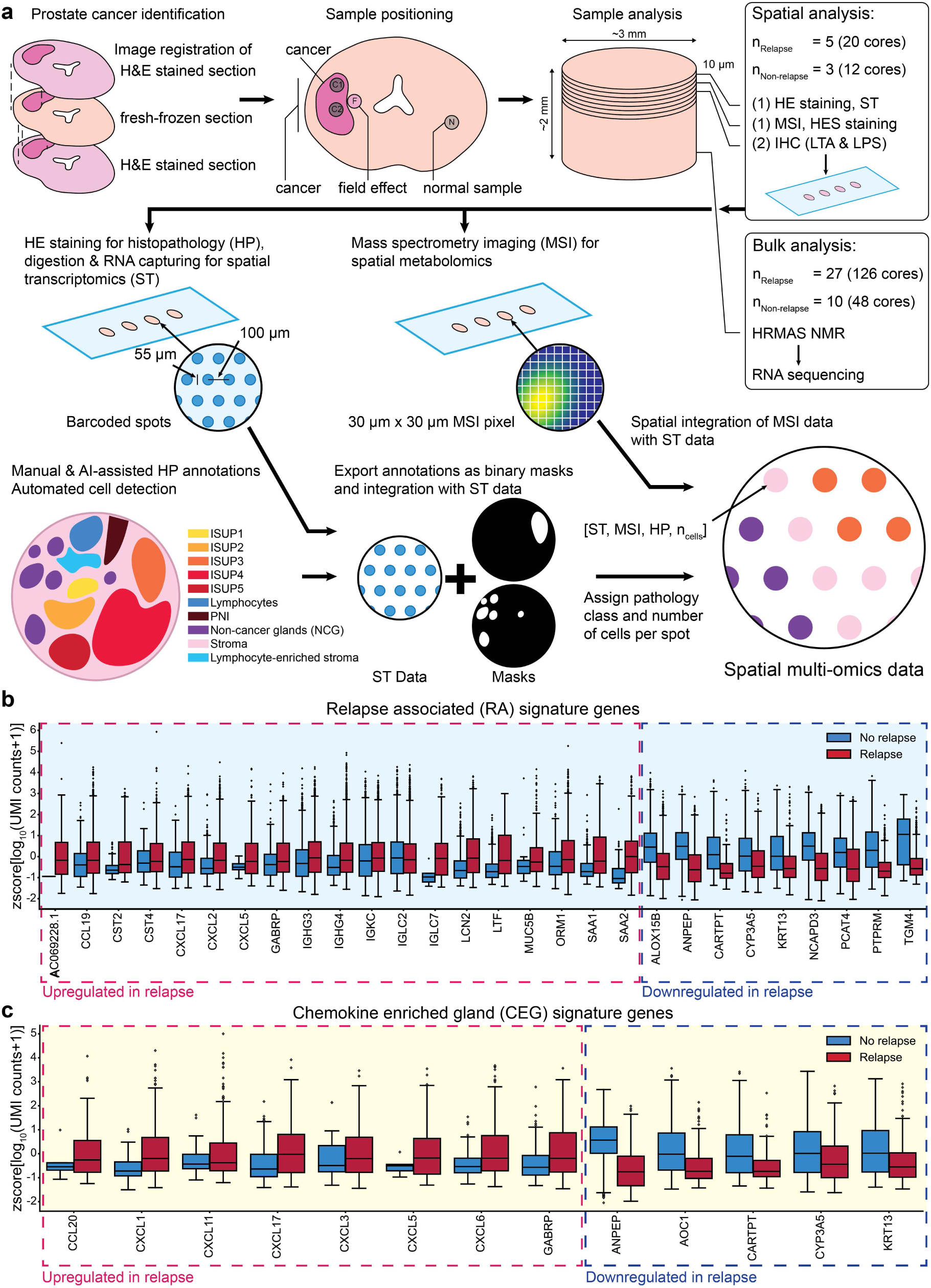
Spatial multi-omics analysis of the prostate tumor microenvironment reveals gene signatures discriminating between relapsing and non-relapsing patients. (a) Schematics of cancer, field effect, and normal sample collection for bulk and spatial analysis of PCa tissue using bulk and spatial transcriptomics (ST), bulk metabolomics (high-resolution magic-angle spinning [HRMAS] NMR), spatial metabolomics (mass spectrometry imaging [MSI]), and histopathology staining (HE[S], immunohistochemistry [IHC] of lipopolysaccharide (LPS) and lipoteichoic acid (LTA)). The fresh frozen prostate tissue sections for sampling were selected based on pathology annotations from adjacent FFPE sections to locate cancer and normal areas. Cryo-sections from samples were obtained at 10 µm thickness and placed on slides for spatial analysis, here exemplary shown for ST and MSI. ST spots were classified according to per-section histopathology and assigned a cell count. Finally, all spatial modalities were integrated into one spatial multi-omics data set. Expression of up- and downregulated genes forming the (b) relapse-associated (RA) signature (c) and chemokine enriched gland (CEG) signature over all spots with a count > 0 for the respective gene separated according to relapse status. The box and whisker plots are based on the z-score of the decadic logarithm of the normalized unique molecular identifier (UMI) counts.

**Table 1.** Clinical data for all patients included in this study. The table is divided by patients of which tissue samples were used for multi-layered spatial ’omics and bulk transcriptomics and metabolomics.

|  | Multi-layered spatial 'omics (n=8) |  | Bulk Transcriptomics & Metabolomics (n=37) |  |
| --- | --- | --- | --- | --- |
|  | Relapse (n=5) | Non-relapse (n=3) | Relapse (n=27) | Non-relapse (n=10) |
| ISUP grade | ISUP2, n=1 | ISUP2, n=2 | ISUP2, n=7 | ISUP2, n=8 |
|  | ISUP3, n=1 | ISUP3, n=1 | ISUP3, n=10 | ISUP3, n=2 |
|  | ISUP4, n=1 |  | ISUP4, n=4 |  |
|  | ISUP5, n=2 |  | ISUP5, n=6 |  |
| Mean age and range at surgery | 59 (53-73) | 60.7 (56-63) | 62.4 (51-73) | 59.8 (55-66) |
| Mean preoperative PSA | 23.7 | 14.0 | 18.7 | 10.8 |
| Clinical T-stage* | T2c, n=1 | T2c, n=2* | T2c, n=9 | T2c, n=9 |
|  | T3a, n=2 |  | T3a, n=6 |  |
|  | T3b, n=2 |  | T3b, n=12 |  |

\*For one patient T-stage information was not available.

### Novel Relapse associated (RA) and Chemokine Enriched Gland (CEG) signatures derived from spatial transcriptomics data

To investigate differences in the transcriptome between relapsing and non-relapsing patients, we first identified sets of differentially expressed genes for each of the histopathology spot classes individually in the ST data. The genes were selected based on highest normalized spatial expression variability and deregulation combined with gene ontology enrichment (GOE)-based evaluation of those genes (**Methods, Supplementary Figure 1a & 2**). The GOE revealed inflammatory and immune-response-related processes as a major difference between these patient groups and evidence hinting at an important role of chemokine-mediated chemotaxis of neutrophils and macrophages (**Supplementary File 2**). We thus decided to emphasize our focus on chemokines by manually including all chemokines detected in the ST data into our signature generation pipeline (see **Methods** and **Supplementary Figure 1b**). We found a set of 26 genes which were able to distinguish relapsed from non-relapsed patients across all histopathology classes (**Supplementary Figure 3**). We termed this set of genes the relapse-associated (RA) signature, which contained 18 genes with higher expression in relapsed patients, and 8 genes with higher expression in non-relapsed patients (**Figure 1b**). Most of the upregulated genes (14 of 18) in the RA signature were directly related to immune response processes. In addition, we observed that the RA signature also scored high in some of the, histologically benign, NCG spots in samples both close and distant to cancer areas from relapse patients (**Figure 2a, Supplementary Figure 1a & 3a**). We therefore derived a gene set particularly capturing the transcriptomic changes in these potentially aberrant NCGs in relapse patients. We termed this the Chemokine Enriched Gland (CEG) signature, reflecting the observed upregulation of 7 chemokines in these benign appearing glands. We calculated for each spot the *single sample Gene Set Enrichment Analysis* (ssGSEA, Markert et al.[24]) scores for the RA and CEG signatures. The RA ssGSEA score showed an enrichment in all histopathology class spots and sample types from relapse patients compared to non-relapse patients (**Figure 2a, b and Supplementary Figure 3a**). Considering the distribution of the CEG signature, the ssGSEA score visualizes its capability to discriminate a sub-group of NCG spots. Although we observed an overall CEG signature enrichment in relapse compared to non-relapse patients, the spot histopathology class with the strongest separation were the NCG spots **(Figure 2c)**. Interestingly, not all NCG spots of relapse patients had a high CEG score, some even had negative scores (compare **Supplementary Figure 1a and 3b** NCG spots at 9 o’clock of section P30 – sample N). In case of high CEG scores, the surrounding stroma was often infiltrated by immune cells as can be seen in sections P08 – sample F & N, P30 – sample C2 (compare **Supplementary Figure 1a and 3b**). To further quantify the histopathological composition captured by both signatures we grouped the ST spots according to the RA and CEG ssGSEA score into groups with significantly high, low, or not significantly changed score (**Supplementary Figure 4**). We found that for both signatures the not significant spot groups had similar compositions of approximately 40% stroma, 23% NCG, 35-40% cancer, and 1% lymphocyte spots. For the high RA score spots, we found a slight increase of stroma spots (56%) and a reduction of cancer spots (18%) while the amount of NCG spots remained stable (22%). The high CEG score spots were composed of an increased amount of NCG spots (55%) and a reduced amount of stroma (26%) and high grade (ISUP >= 3, PNI) cancer spots (3%). In contrast, the low score groups for both signatures consisted nearly completely of NCG spots (RA 80%, CEG 95%). In conclusion, we found two novel ST signatures with the RA signature displaying relapse-associated transcriptomic changes independently of the histopathology while the CEG signature was characteristic for NCG and low-grade cancer spots of relapse patients.

**Figure 2.**
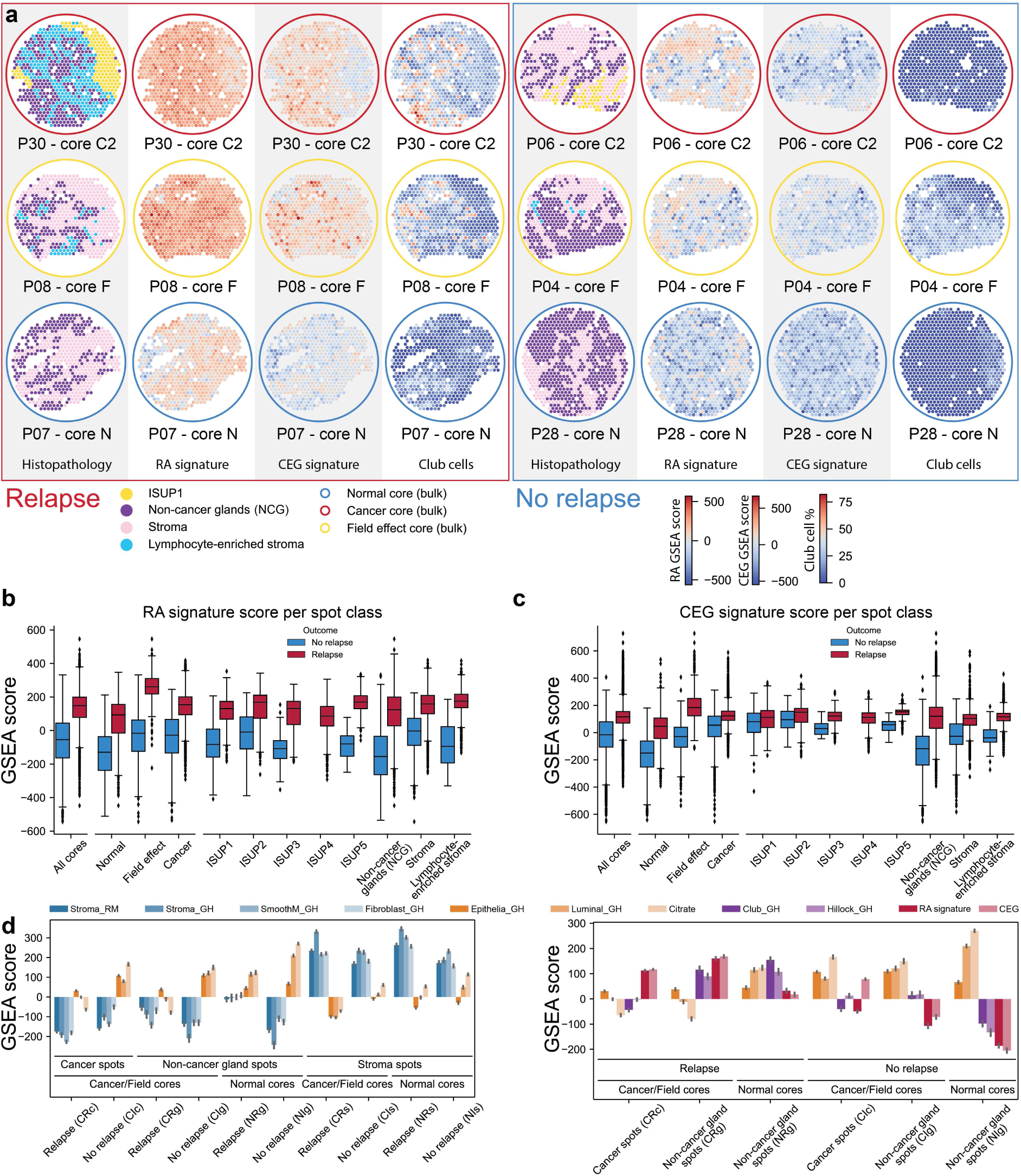
RA and CEG signature activity in prostate tissue. (a) Spatial distribution of RA and CEG signature activity, exemplary shown for one sample type for each patient group, visualizes the global and uniform activity of the RA signature as compared to the CEG signature. Spots with high CEG signature activity contained high estimated Club cell fractions. (b) RA signature activity was higher in all spot histology classes from relapsing patients (red) compared to non-relapse (blue), while (c) CEG signature activity was higher in non-cancer gland (NCG) spots and less pronounced in stroma and cancer spots of relapsing patients and more prevalent in normal and field effect samples. (d) Mean scores of stromal activity gene sets (blue) in spots grouped by histopathology class (cancer, non-cancer gland, stroma), sample type (cancer & field, normal), and relapse status. Stromal activity was increased in stroma spots and decreased in NCG spots and vice-versa for selected epithelial gene sets (orange). (e) Mean activity of selected prostate epithelial gene sets, cell type specific gene set, RA, and CEG signature showed clear differences between relapse and non-relapsing patients. Both, CEG and RA signature showed increased activity in relapsing patients accompanied by an increased activity of the Club signature and a slight reduction, especially in the NCG spots close to cancer, in luminal and citrate/spermine gene sets.

### RA and CEG signature are associated with loss of luminal features and citrate secretion in glands

Considering that both our gene signatures contained a high number of immune-related genes, we investigated potential associations with molecular processes and cell-types. Further, based on the observation that both signatures highlighted a subset of high scoring NCG spots, we investigated if these spots, despite their benign appearing histology, already underwent molecular aberrations associated with cancer. Therefore, we calculated for each spot ssGSEA scores using 38 gene sets characteristic for cell types and cellular phenotypes related to either normal prostate or PCa tissue (**Supplementary Figure 5 and Supplementary Table 2**). We suspected the distance to cancer could influence such changes and thus defined 10 pseudo-bulk groups of ST spots based on the three major histology classes (cancer, NCG, and stroma), sample type (cancer, field effect, and normal) and patient relapse status to simplify the analysis and still preserve to some degree the spatial nature of the data. Because of the low number of field effect samples, we merged cancer and field effect samples unifying spots close to cancer and to obtain reasonable spot numbers in each of the 10 pseudo-bulk groups (863 to 4263 spots per group) to allow a reliable ssGSEA. We first validated our ssGSEA approach using gene sets for stroma and gland, the two most well-defined prostate tissue compartments [23]. All stroma spot groups showed strong enrichment for the four stroma-related signatures (**Figure 2d**). The gland-related gene signatures ‘Epithelia_GH’, ‘Luminal_GH’, and ‘Citrate’ (see **Supplementary Table 2** for details) were generally enriched in normal sample NCG spot groups with higher scores in spots from non-relapse patients (**Figure 2d**), confirming the validity of our pseudo-bulk approach for ST data. Further, we observed that these gland signatures were depleted in NCG spots close to cancer in relapse patients but not in non-relapsing patients. Interestingly, CEG and RA signatures followed an inverse trend, they were enriched in relapse patients where the gland signatures were depleted and vice-versa (**Figure 2e**). Additionally, we observed normal gland signature enrichment in cancer spots from non-relapse patients, but depletion in relapse patients. In conclusion, we found that high activity of both signatures in benign appearing glands was associated with transcriptional loss of luminal features and citrate secretion.

### RA and CEG signature are associated with Club-like cell and immune cell enrichment

Considering the observed associations of the RA and CEG signatures with reduced luminal characteristics we wanted to gain a better understanding of underlying processes by looking at potential associations with specific cell populations present in the tissue of relapse patients. Thus, we first investigated which of the cell-type specific gene sets of the 38 used in the pseudo-bulk ssGSEA approach were co-enriched in the same tissue regions, specifically in NCG spots close to the cancer (**Supplemental Figure 5b**, CRg group). We saw an enrichment of immune cells from the lymphoid and myeloid lineages in this spot group (CRg) and even in the NCGs further away from the cancer (NRg) of relapse patients compared to the respective groups (CIg & NIg) in non-relapsing patients (**Supplemental Figure 5b**). Of note, we also observed an enrichment of the gene sets for Club and Hillock cells in these spot groups from relapsing but not from non-relapsing patients (**Figure 2e and Supplemental Figure 5b**).

Motivated by the correlation of our signatures with these cell type specific signatures, we further investigated signature and cell-type correlations on the individual spot level. Since ST data captures RNA from roughly 10-30 cells in each individual spot, the spot expression profiles originate from a mixture of cell-types. Thus, we choose to deconvolute each spot using *stereoscope*, a tool specifically developed to estimate cell-type compositions in ST spots by learning cell-type specific gene expression profiles from single-cell RNA-sequencing data [25]. The stereoscope model was trained on publicly available single-cell RNA-sequencing data obtained from normal prostate tissue [10].

Using this approach, we found that the overall cell-type distribution was dominated by epithelial cells, followed by a large fraction of smooth muscle cells, a comparably large fraction of immune cells, and smaller fractions of fibroblasts and endothelial cells (**Figure 3a**). Interestingly, the most prominent difference between relapse and non-relapse spots was the relative increase of immune cells and Club-like cells (**Figure 3a**). The correlation of histopathology spot classes with cell-type composition showed that NCG spots correlated positively with epithelial cell types and negatively with fibroblasts and smooth muscle cells while stroma spots inversely correlated with these cell-types **(Supplementary Figure 6)**. Further, luminal epithelial cells were only weakly positively correlated with ISUP1 spots, and this correlation gradually decreased with increasing ISUP grade to reach a weak negative correlation with ISUP5 spots, reflecting the loss of luminal characteristics in this tissue. These correlations of cell types and histology are comparable to our observations using the ssGSEA pseudo-bulk approach and thus validated the underlying stereoscope model.

**Figure 3.**
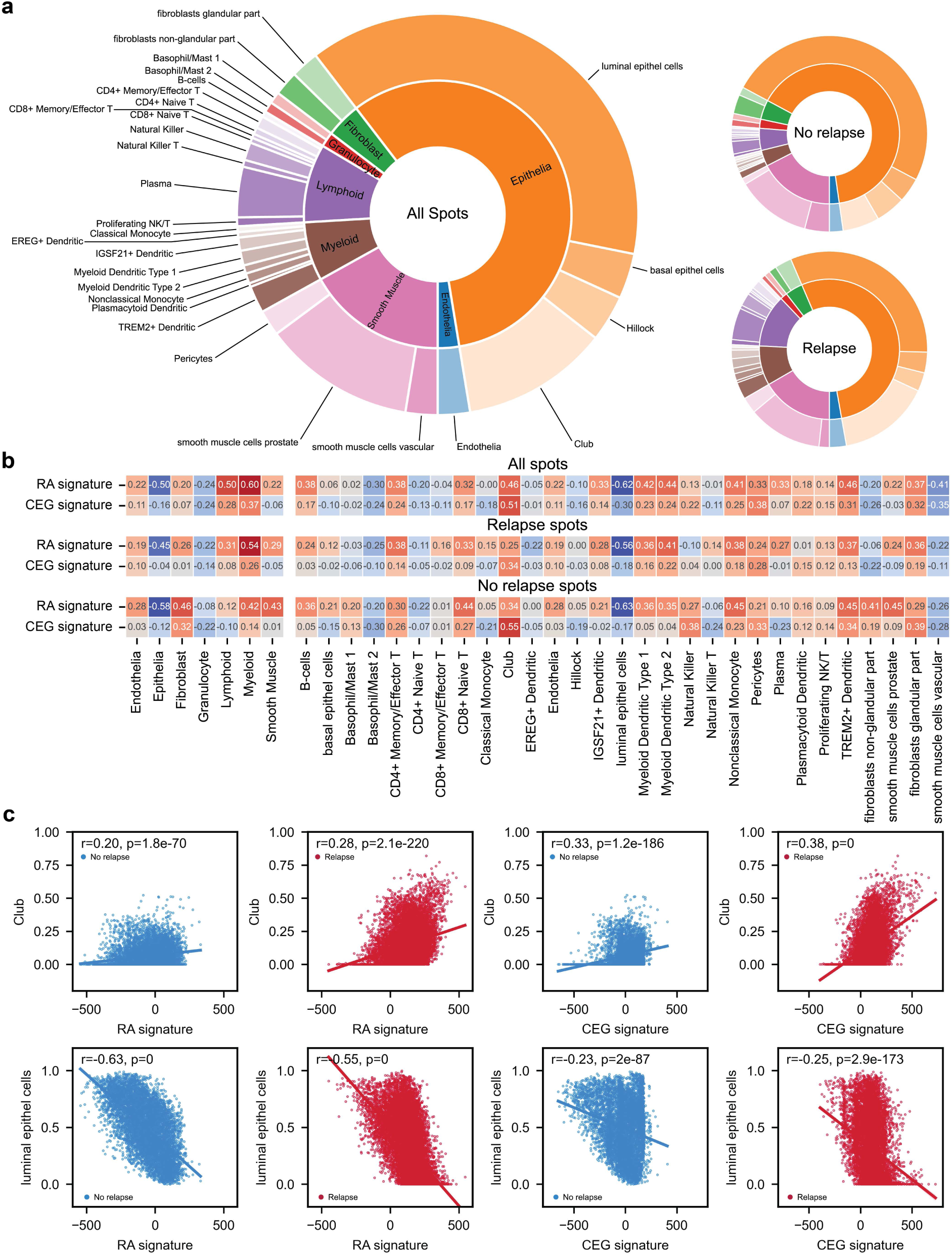
Cell-type composition of each ST spot and correlation with RA and CEG signatures. Using cell-type specific RNA expression profiles derived from single-cell RNA-sequencing data we were able to deconvolute each ST spot into its cell-type composition. (a) The accumulated cell-type fractions are shown for all spots (left) and separated by relapse status (right). (b) Spearman correlation coefficients calculated for cell-types and signature activity for all spots (top), spots from relapsing patients (middle) and spots from non-relapsing patients (bottom) visualized by color (red = positive, blue = negative correlation). (c) Scatterplots with linear regression of RA and CEG signature ssGSEA score vs. Club and luminal epithel cells contents individually for spots from relapsing (red) and non-relapsing (blue) patients. The R- and p-values for each linear regression are given inside the box of each plot.

To further narrow down cell type associations with the RA and CEG signatures, we calculated the spearman correlation of the signatures with the stereoscope-derived cell fractions. We found that immune cells of the lymphoid and myeloid lineages, but not granulocytes, were positively associated with the RA and CEG signatures. Further, the strongest negative correlations were observed for RA and CEG signatures with luminal epithelial cells and the strongest positive correlations with Club-like cells **(Figure 3b, c)**. No clear correlation with hillock cells was observed **(Figure 3b)**. The correlation with Club-like cells agrees with our observation from the ssGSEA pseudo-bulk analysis, while for the hillock cells there were conflicting results. The KRT13 gene is downregulated in both signature active regions (**Figure 1b**) and hillock cells are characterized as KRT13+. Hence, the weak negative correlation of the stereoscope-derived hillock cell fractions with the signatures might reflect this. Nevertheless, Hillock cells are not well described in the literature, especially in the human prostate, making it likely that both approaches (stereoscope and ssGSEA) might capture other closely related cell types. Thus, we decided to focus on the enrichment of Club-like cells in NCG spots close to the cancer of relapsing patients that also had a depletion of luminal and citrate secretion signatures. Overall, loss of luminal characteristics and lower citrate secretion in these NCG spots indicated a switch from a healthy prostate gland phenotype to the molecular phenotype of PCa. In conclusion, these “defunctionalizing” NCG spots in relapse patients were enriched with Club-like and immune cells while simultaneously characterized by high RA and CEG signature activity.

### Chemokines in RA and CEG signature likely to originate from Club-like cells

To gain further insight into the cellular origin of the individual genes of our signatures, we looked at the correlation of those genes with the estimated cell type compositions (**Supplementary Figure 7**). For the correlation of upregulated RA signature genes with Club-like cells we found clear positive correlation (ρ >= 0.25) for the genes LCN2, LTF, SAA1, SAA2, CXCL17 and weak positive correlation (ρ >= 0.15) for the genes GABRP, IGHG3, IHGH4, IGLC7, MUC5B, CCL19, CXCL2, CXCL5 (**Supplementary Figure 7a**). Cells of myeloid lineage representing the first line of defense against infections were also positively correlated with these genes and further strongly (ρ >= 0.37) positively correlated with LTF, CCL19, and some of the immunoglobulins upregulated in the RA signature (IGHG3, IGKC, IGLC2). The same immunoglobulins in addition to IGHG4 were also strongly positively correlated with lymphoid cells. In contrast, luminal epithelial cells were positively correlated with the downregulated genes ALOX15B, ANPEP, CARTPT, CYP3A5, NCAPD3, PCAT4, and PTPRM (ρ >= 0.20). For the CEG signature genes (**Supplementary Figure 7b**), we found that the additional chemokines CXCL1, -3, -6, -11, and CCL20 were all also positively correlated (ρ >= 0.10) with Club-like cells and cells of the myeloid lineage. Of note, the expression of these genes, apart from CXCL11, was mostly negatively correlated with luminal epithelial cells and other cell types of the lymphoid or endothelial lineages. Considering that Club-like cells are known to secret chemokines [14], these correlations suggest Club-like cells as the likely source for the chemokines of our signatures.

### Atypical Chemokine Receptor 1 (ACKR1) is the dominant receptor for CEG-signature CXC-chemokines in relapse patients

Chemokines interact with specific chemokine receptors typically resulting in chemotaxis of the target cells. After identifying Club-like cells as the likely source for the chemokine secretion captured in the CEG signature, we investigated which chemokine receptors were detected in the ST data and what potential receptor-ligand interactions were active. Of all the chemokine receptors, we only detected expression of CXCR4 and ACKR1 in the ST data (**Supplementary Figure 8a**) of which only ACKR1 is known to respond to the CXC chemokines of the CEG signature. Bulk transcriptomics data obtained from 174 samples of 37 patients (including the 8 patients of the ST data cohort) demonstrated that both CXCR4 and ACKR1 together with ACKR3 and CCR1 are amongst the highest expressed chemokine receptors in prostate tissue (**Supplementary Figure 8b**). Nevertheless, to investigate potential signaling across different tissue types of the chemokines of the CEG signature in relapse patients, we performed a cellPhoneDB analysis of the ST data using the histopathology classes as groups. To avoid a potential bias caused by data sparsity but also preserve some of the spatial information we performed the analysis for all the relapse samples individually and combined the results but abstained from any further grouping using spot-to-spot distance or cell-type composition.

We found significant interaction of CXCL1, -3, -5, -6, and -11 with ACKR1 from NCG to lymphocyte, stroma, and lymphocyte-enriched stroma spots (**Figure 4a**). Corresponding interactions were also observed from stroma and lymphocyte-enriched stroma to lymphocyte, stroma, and lymphocyte-enriched stroma spots. Interestingly, CXCL1, 3, and 6 – ACKR1 interactions were also present with NCG spots as origin and ISUP1 and NCG spots as receiver. Of note, we found a CXCL11-DPP4 interaction with NCG, stroma, and lymphocyte-enriched stroma spots as sender and NCG and cancer spots as receiver. The expression level of CXCL1, -3, -5, -6, -11, ACKR1, and DPP4 in spots from relapsing patients supported these interactions (**Figure 4b**). We found NCG and stroma spots to be the main source for these CXC chemokines while for the cancer spots the expression was lower and mainly detected in the low-grade spots. ACKR1 was detected in all spot classes but was highest in both stroma classes and the lymphocyte spots. Compared to ACKR1, DPP4 showed a high expression in all spot classes except for lymphocyte spots. To gain insight into the cell types that express ACKR1 and DPP4, but also CXCR4, we looked at the correlation between their expression and the cell type composition for each ST spot (**Figure 4c**). ACKR1 was positively correlated with endothelial cells (ρ = 0.29) and slightly negatively correlated with luminal epithelial cells (ρ = -0.12). In contrast, DPP4 was negatively (ρ = -0.20) correlated with endothelial cells and strongly positively (ρ = 0.53) correlated with luminal epithelial cells. CXCR4 was positively correlated with cells of the myeloid (ρ = 0.37) and lymphoid (ρ = 0.28) linages and, like ACKR1, also negatively correlated with luminal epithelial cells (ρ = -0.26). To conclude, we found ACKR1 to be the dominant receptor for CXC chemokines in high CEG signature scoring spots and likely expressed by endothelial cells.

**Figure 4.**
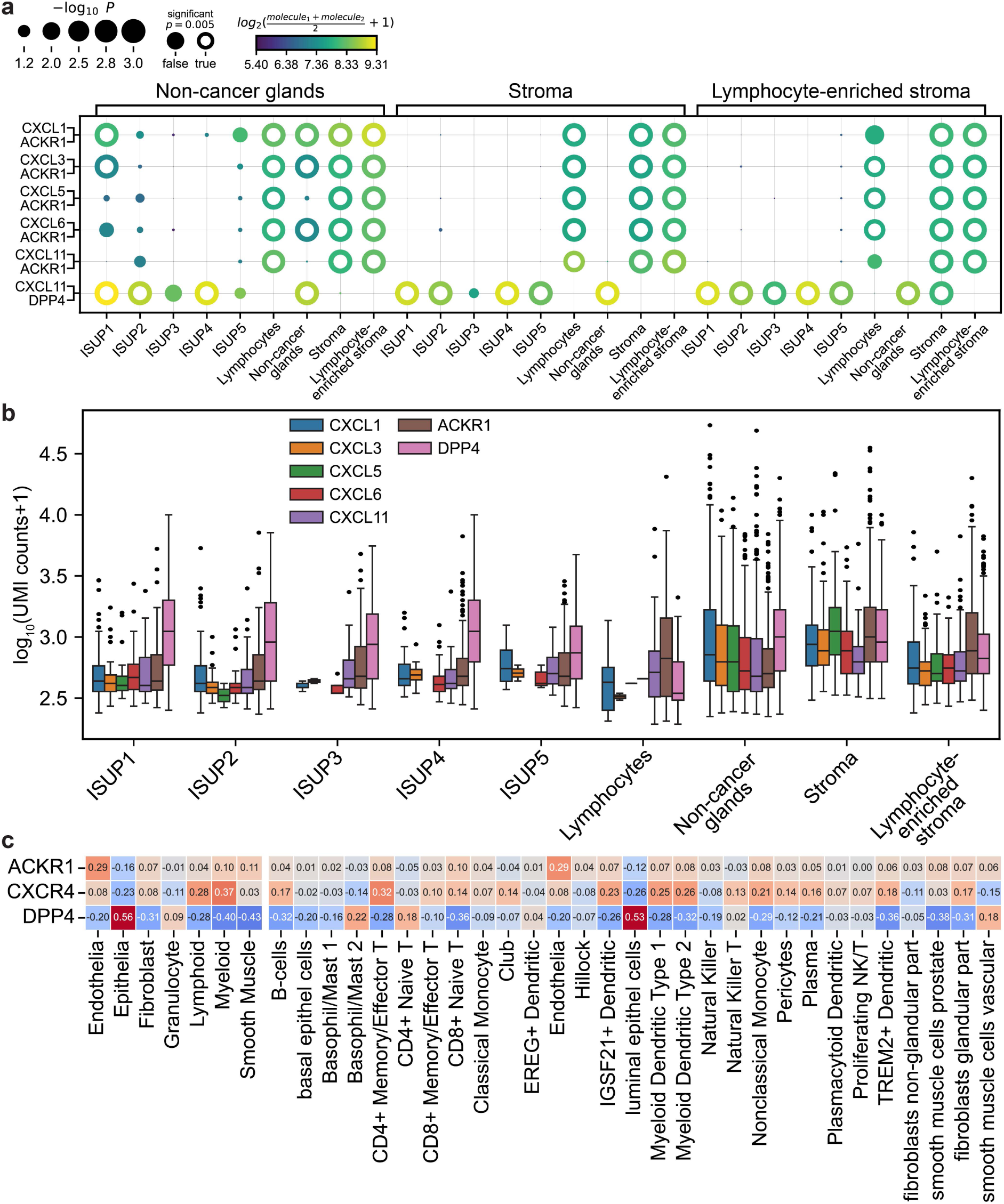
Chemokine receptor-ligand interaction in tissue from relapse patients. (a) Potential receptor-ligand interactions of the CXC chemokines of the CEG signature with ACKR1 and DPP4 between different tissue types found by combining the CellPhoneDB results obtained from ST data of each of the samples from relapsing patients. The means of the average expression of the receptor-ligand pair are visualized with a color gradient and the p-values by the size of the circles. Receptor-ligand pairs are shown on the y-axis and interacting spot class pairs are given on the top and bottom x-axis. (b) Expression of the CXC chemokines of the CEG signature, the chemokine receptor ACKR1, and DPP4 in spots with UMI count > 0 from relapse patient tissue. (c) Correlation of ACKR1, CXCR4, and DPP4 expression with cell type fractions per spot, colored according to Spearman correlation coefficient (blue = negative, red = positive correlation). Correlations for the CXC-chemokines are visualized in Supplemental Figure 6.

### No obvious correlation of bacterial infections and CEG signature

Acute and chronic bacterial prostatitis are common [26] and thus a likely cause for the inflammation we observed in our tissue samples. Despite this, we could not detect expression of cytokines typically induced by bacteria during acute infection, such as IL-6 and TNF-α in the ST data. We further investigated this by staining serial sections for lipopolysaccharides (LPS) and lipoteichoic acid (LTA) for the detection of gram-negative and gram-positive bacteria. Following affine co-registration of the serial sections we calculated an interpolated RA and CEG signature activity density map for each of these stained sections. Next, we ran an automated cell detection and assigned an estimated signature activity and staining intensity to each cell (**Supplementary Figure 9**). Only a small fraction of the cells stained positive for LTA or LPS (359 of 718 901 and 374 of 746 976 cells, respectively) and we did not find an association of RA or CEG signature activity with LTA or LPS staining. To conclude, it is unlikely that an active infection is the cause for the observed inflammation.

### RA and CEG signature activity increased in bulk samples from relapse patients

The highly detailed, spatially resolved data from 8 patients allowed us to find both signatures and characterize associated biological processes. Unfortunately, in day-to-day clinical practice such data is rarely available and such detailed analysis hardly feasible. However, prostate biopsies are typically collected and easier available for bulk molecular diagnostics. Thus, we wanted to elucidate if the RA signature and the CEG signature were detectable, predictive of relapse in bulk samples of a larger patient cohort. We collected and analyzed RNA-sequencing data of 174 (no relapse n=48, relapse n=126) samples (classified as either cancer, field effect, or normal) from 37 (no relapse n=10, relapse n=27) patients (**Supplementary Figure 10 & 11**). The ssGSEA scores for the RA signature were higher in samples from relapsing patients (**Supplementary Figure 10a**). The strongest signal originated from cancer samples while the difference in normal and field effect samples was less prominent. Nevertheless, receiver operating characteristics (ROC) calculated using the median signature score per patient resulted in an area under the curve (AUC) of 0.75 (**Supplementary Figure 10b**). A per patient analysis revealed strong patient to patient variations with at least one clear outlier in the non-relapsing group (P05, **Supplementary Figure 10c**). Further, for the CEG signature we detected a weaker difference in activity between relapse and no relapse patients as compared to the RA signature with an AUC of 0.66 of the ROC (**Supplementary Figure 11**). Both groups had samples with high and low signature activity to the point that on average the signature activity was even slightly higher in normal and field effect samples of non-relapsing patients. A closer look at the per patient signature activity distribution (**Supplementary Figure 11c**) revealed that this effect was mainly caused by patient-to-patient variability. For example, non-relapsing patients P05 and P32 had samples with high signature activity while relapsing patients P09 and P25 had low signature activity. Interestingly, the overall difference – the variation around the median activity – was also much smaller in bulk data as compared to ST data.

### RA and CEG signature associated with altered metabolism

After extensive and in-depth analysis of spatial and bulk transcriptomics data we investigated if the loss of luminal features and citrate secretion on the transcriptional level in RA and especially CEG high scoring ST spots resulted in a measurable change of related metabolites. Further, we hypothesized that the inflammatory processes captured with our signatures were likely chronic and we investigated if they are correlated to metabolic aberrations. To reveal such changes, we examined spatial metabolomics data obtained by MALDI MSI on serial sections adjacent to sections that were used for ST (**Figure 1a**). MSI data were registered to the corresponding ST data and median relative levels of putatively identified metabolites for each ST spot were calculated using our Multi-Omics Imaging Integration Toolset (MIIT) [27]. To analyze potential metabolic changes associated with RA and CEG signature activity, we applied the same grouping of spots into significantly high, low, or not significantly changed RA and CEG score as initially used for quantifying the histopathology composition captured by our signatures. Based on the composition of these groups, we considered the metabolite composition of the low signature score groups as most likely resembling healthy normal prostate glands while the not significant spot groups captured an average PCa tissue metabolic state. Consequently, we found that the signature low scoring spots had on average the highest levels of citrate and zinc while the not significant scoring spots had reduced levels of both (**Figure 5**). For the high scoring spots this reduction was even more pronounced. Further, even though less pronounced, we found that ATP levels were also highest in low scoring spots as compared to not significant and high scoring spots. In contrast, we detected an inverse trend for AMP, ADP, glutathione, glucose, glutamate, glutamine, *N*-acetylaspartate (NAA), urate, and taurine. For all these metabolites we detected the lowest levels in low scoring spots and highest levels in high scoring spots. Aspartate was the only detected metabolite with similar levels in all spots. Interestingly, we detected the same trends and similar average relative metabolite levels for both RA and CEG signature high scoring spots despite their different histopathology compositions. Next, we analyzed the obtained metabolite data from our bulk samples by grouping, correspondingly to the spatially resolved data, by significantly high, low or not significantly changed signature activity (**Figure 5**). We observed a comparable trend for citrate with the highest levels detected in low scoring samples and lowest detected in high scoring samples. Correspondingly, we found glutamate and taurine levels to be lowest in low scoring samples and highest in high scoring samples. Compared to spatial data these trends were less pronounced. Further, we found that lactate levels were slightly lower in low scoring samples as compared to not significant and high scoring samples. Interestingly, we found succinate to be the only metabolite with different levels between RA and CEG high scoring samples. Grouping bulk samples by RA signature score did not display any trend while for CEG the low scoring samples showed comparably lower succinate levels. In conclusion, we found clear indications for metabolic alterations in high RA and/or CEG scoring spots in agreement with transcriptional loss of luminal functions and citrate secretions.

**Figure 5.**
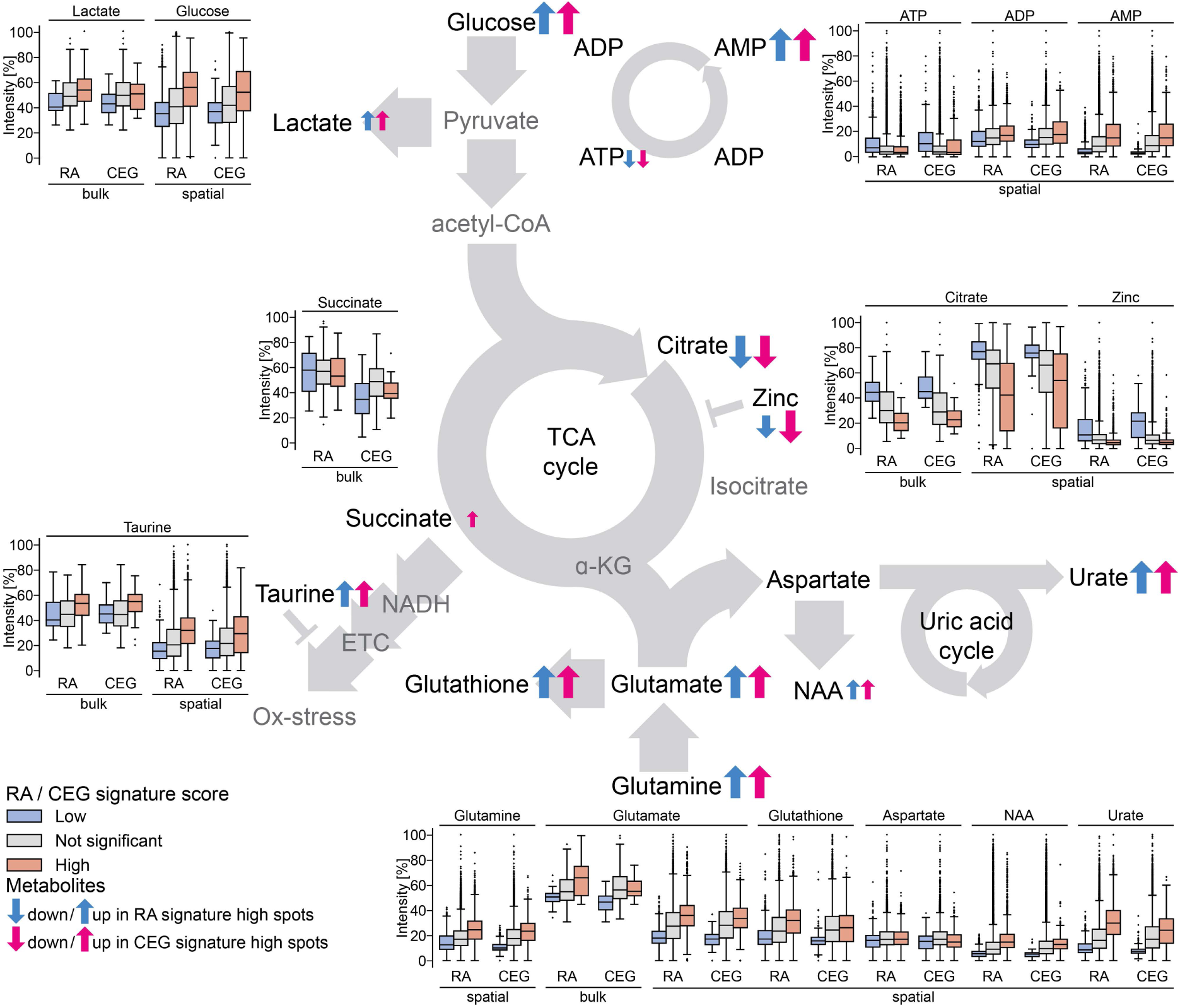
Changes of metabolite levels in relation to RA and CEG signature activity. Shown are the relative intensities in % grouped by signature activity of metabolites detected either by MSI (“spatial”) or HRMAS NMR (“bulk”) placed according to their position in the indicated simplified metabolic pathways (glycolysis, TCA cycle, uric acid cycle). Metabolite level changes (up or down) in spots (spatial) or samples (bulk) with an increased signature activity compared to low signature activity are indicated with up or down arrows (blue = RA, magenta = CEG signature)..

### Analysis in public bulk data supports association of RA and CEG signature with Club-like cells and reduced luminal function

To investigate generalizability of our results, we analyzed bulk gene expression data from 12 publicly available PCa cohorts comprising a total of 2512 tissue samples (**Supplementary Table 3**). We calculated Pearson correlations between ssGSEA scores of RA, CEG, and Club gene set-based signatures with scores for the gene sets used previously in the ST data (**Supplementary Figure 5**). To estimate the correlation of our signatures with aggressive PCa, we also included gene sets based on three commercially used gene expression signatures (CCP, Decipher, GPS) for aggressive PCa in this analysis. Correlation analysis in public data in general confirmed the associations observed in ST data (**Figure 6, Supplementary File 3**)

**Figure 6.**
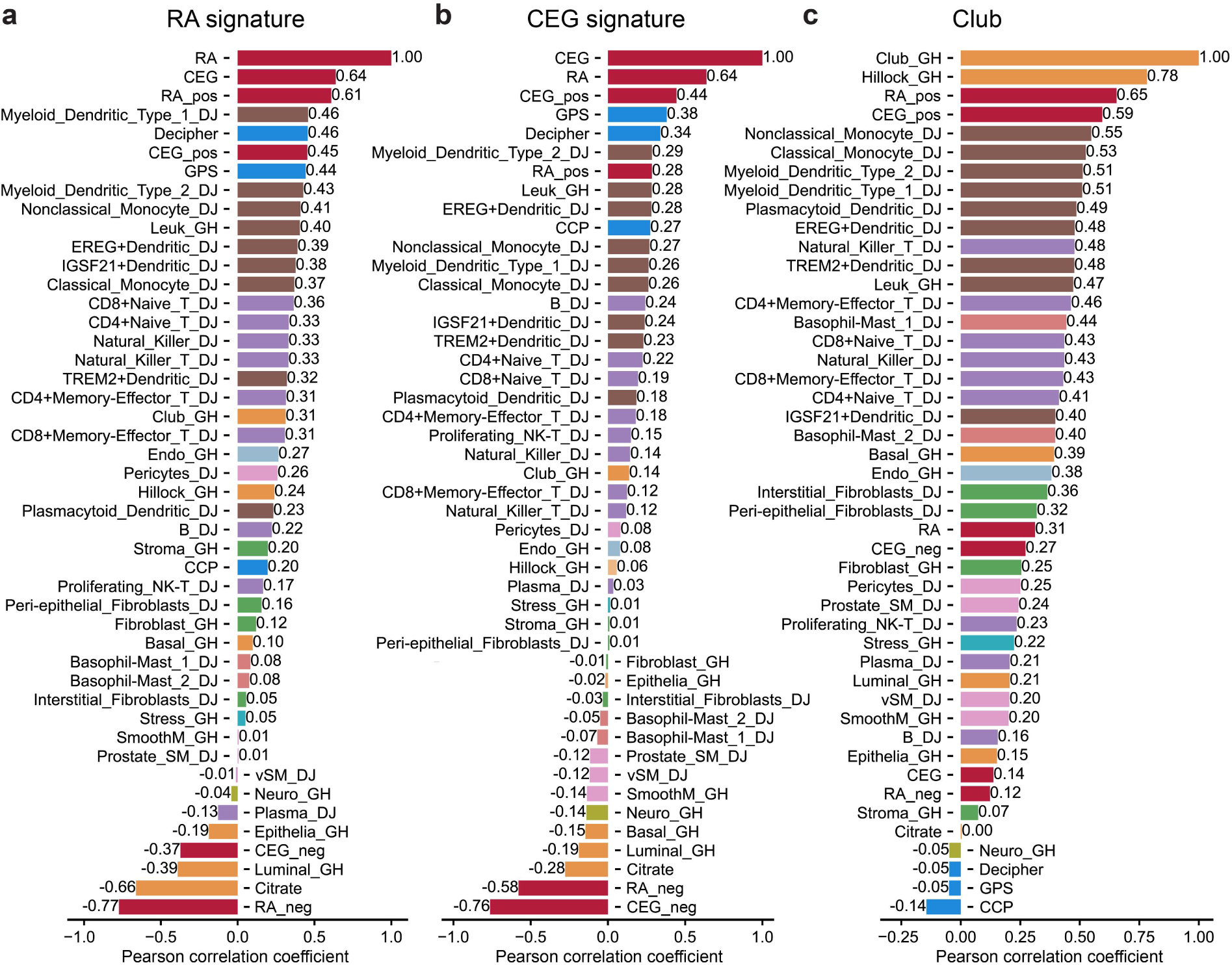
Average Pearson correlation between gene sets across 12 public data sets for a) RA, b) CEG and c) Club cell signature. Correlations are displayed for RA and CEG signature with positive (upregulated genes) and negative (downregulated genes) counterparts (dark red), epithelial linage gene sets, like Club, Luminal cells, and Citrate gene sets (orange), immune cell gene sets (lymphoid –purple, myeloid –brown, granulocyte – light red), and genes from signatures known to be associated with aggressive PCa (GPS, Decipher and CCP, blue). Correlations for all gene sets and individual public data sets are shown in Supplementary File 2 – Public Data.

The RA signature correlated with genes from GPS (ρ=0.44) and Decipher (ρ=0.46), which supports the association between the RA signature and aggressive, recurrent cancer. The correlation was less strong for the CCP genes. The CEG signature also showed similar correlation preferences; however, the correlation values were weaker than for the RA signature. Both signatures also showed positive correlations to immune cells, but also here the correlation values were weaker for the CEG signature. Both signatures correlated negatively with the Citrate signature. Interestingly, the negative correlation to the Citrate signature was stronger compared to the negative correlation of Luminal cells, highlighting the relation of both our signatures with the secretory function of the luminal cells.

The RA and CEG signatures showed only intermediate (RA) or weak (CEG) positive correlation to Club-like cells in the public datasets. However, when we separated both signatures into their positive and negative contributions (up- and down-regulated genes respectively), we observed strong correlation with Club-like cells for the positive parts of both signatures (**Figure 6c**). This confirms that the observed associations between the two signatures with Club-like cells are extendable to larger patient cohorts. Interestingly, though inversely correlated in the ST data, the positive and negative counterparts of the CEG-signatures were not inversely but slightly positively correlated in the bulk data (**Supplementary File 3**). This could indicate that we are able to observe local compartmentalized signature combinations in ST data that is not detectable in bulk data.

## Discussion

Finding good biomarkers for PCa that can predict aggressive disease progression early is a challenging task and precise analysis of the relation of potential markers with known and potential new functional aspects of PCa is essential. Using ST data, we could identify two signatures of gene expression markers capturing changes in the TME of PCa and link them to biological functions and metabolic changes.

Both signatures mainly captured upregulated inflammation related signals highlighting the role of an inflammatory immune response in aggressive PCa. Specifically, the RA signature (**Figure 1 b**) contained the acute-phase proteins serum amyloid A (SAA1 and SAA2) and alpha-1-acid glycoprotein 1 (ORM1) as well as immune globulins (IGHG3, IGHG4, IGKC, IGLC7). The immune globulins can be seen as a direct gene expression-based indicator for the histologically observed increased presence of immune cells in tissue of relapsing patients. While serum amyloid A and alpha-1-acid glycoprotein 1 can be interpreted as markers for acute and chronic inflammation they have also been implicated in tumor pathogenesis and have been suggested as serum-based biomarkers for aggressive PCa and biochemical recurrence, respectively [28-30]. Cystatin SA (CST2) and Cystatin S (CST4) are known to be present in seminal fluid, have anti-viral properties and elevated levels have been reported for many cancer types including PCa [31-33]. Of note, we found lactotransferrin (LTF) upregulation to be associated with relapse in our data. While LTF has been reported as a tumor suppressor and immune modulator in PCa [34], a recent study in glioblastoma multiforme demonstrated that high LTF expression was associated with worse survival and increased immune cell infiltration [35]. Lipocalin-2 (LCN2) is known to be expressed and TNF-ɑ-inducible in the prostate and a tumor and proliferation promoting factor [36, 37]. Further, LCN2 together with the chemokine CXCL1 were reported as prognostic markers for PCa relapse and facilitated metastasis in a mouse model [38]. The role of the mucine MUC5B in PCa has not been studied much but there are results implicating a role in hormonal escape of PCa [39]. Similarly, GARBP, the π-subunit of the GABA A receptor, has only been shown to be involved in in vitro PCa cell line proliferation [40]. Interestingly, our RA signature also contained the long non-coding RNAs AC069228.1 and PCAT4. Their distinct role in PCa remains to be investigated, but two studies linked AC069228.1 to relapse [41, 42]. Reduction of ALOX15B, PTPRM, and ANPEP have been reported as progressors of and were associated with high-grade PCa [43-46]. Interestingly, NCAPD3 has supposedly tumor-promoting effects in PCa [47] while expression was also associated with reduced recurrence after radical prostatectomy [48], thus confirming our observation that a reduction was associated with more aggressive PCa. Further, CYP3A5 is known to be a modulator of androgen receptor signaling in PCa [49] while the role of CARTPT, a known oncogene, in PCa is not well understood [50]. In contrast, keratin 13 (KRT13) has been suggested as a marker for bone metastasizing PCa [51], while we found downregulation in PCa tissue to be associated with PCa relapse. KRT13 is also a hillock cell marker and appears to play a role in the development of the prostate but further research is required to uncover all functional aspects [52]. Downregulation of SYCE1L has not been reported as marker of relapse or aggressive PCa before. The signature genes LCN2, LTF, MUC5B, CXCL1, CXCL6, and CXCL17 are enriched in so called effector Club cells of the lung tissue playing a crucial role in the first line of immune response [14]. Our findings indicate that the observed enrichment of the chemokines CXCL1, 3, 5, 6, 11, 17, CCL20 in the CEG signature was probably caused by a population of Club-like cells similar in function to lung effector Club cells in the respective NCGs causing the observed infiltration of immune cells potentially amplifying the chemokine signaling. Except for CXCL17, all of these chemokines are pro-inflammatory and attract immune cells [53] and are implicated in pro- or anti-tumor immunity depending on the respective chemokine ligand-receptor interaction and immune cell type [54].

Even though the RA signature was composed of many innate immune response factors and bacterial prostatitis has a high incidence [26], using IHC, we did not detect any signs of an active bacterial prostatitis as cause for the observed inflammatory response in the tissue. Regarding chronic bacterial prostatitis as a contributing factor, the results remain inconclusive. The detected low-level presence of LPS and LTA in regions with CEG signature activity might suggest some bacterial contribution possibly at an earlier time-point. The individual genes present in both the RA and CEG signature and their correlation with Club-like cells provide evidence of potential pro-inflammatory chemokine signaling from these prostate effector Club-like cells and a potential role in aggressive PCa. The association of Club-like cells with our signatures was further supported by using independent publicly available data.

Our analysis of potential chemokine receptor-ligand interactions in the TME of PCa revealed CXCR4 and ACKR1 as the main receivers of CXC-signaling of which only ACKR1 is known to bind the CXC-chemokines of our signatures. ACKR1, also known as the Duffy antigen receptor, was initially discovered on erythrocytes and loss of expression is associated with malaria resistance [55]. Apart from erythrocytes, ACKR1 is expressed on endothelial cells of the lungs. Here ACKR1 is involved in propagation of CXC chemokine signaling in inflammatory processes, for example by translocating chemokines from the basal to the apical side resulting in the recruitment of circulating immune cells. Considering that our data suggests ACKR1 to be mainly expressed on endothelium in the prostate and the observed increased immune cell infiltration a similar role in the prostate as in the lungs appears plausible. Interestingly, ACKR1’s role in cancer has been reported as dominantly protective against tumor metastasis and uncontrolled proliferation, and is associated with positive outcomes [56]. Nevertheless, in PCa, reduced ACRK1 expression on erythrocytes in men of African descent appeared not to be linked to aggressiveness [57, 58] while ACKR1-deficient mice showed increased tumor growth [59]. In contrast, ACKR3 expression together with CXCR4 is associated with cancer promoting effects [60]. Even though, we only detected ACKR1 and CXCR4 in the ST data of those receptors, we found ACKR3 to be equally high expressed as ACKR1 in our bulk transcriptomics data. DPP4 is known to cleave chemokines at the N-terminus resulting in altered signaling and diverse effects on inflammatory processes [61]. Considering that we found ANPEP, another peptidase known to correspondingly modulate CXCL11 signaling [62], to be downregulated in relapse patients, puts the proposed CXCL11-DDP4 interaction into an interesting perspective strongly suggesting further studies on the role of peptidase-based alteration of chemokine signaling in PCa.

We investigated biological functions associated with our signatures by looking at the activity of known prostate function-related gene sets in our ST data (Figure 2). Our results support our previously reported observations that prostate secretory genes are downregulated in tumors from relapse patients [44]. We could further substantiate this observation by analyzing metabolite compositions in RA and CEG active regions. Our findings show that high activity of both signatures is associated with metabolic changes in the respective regions. Specifically, the metabolic phenotype observed in RA high scoring spots displayed the typical hallmarks of PCa metabolism: Reduced citrate and zinc, increased levels of glucose, and signs for an increased glutamine consumption [63]. Interestingly, the high CEG signature regions, even though mainly containing NCG spots, showed a very similar, cancer-like, metabolic phenotype (**Figure 5**). Zinc is typically understood to block aconitase to allow healthy prostate glands to accumulate high levels of citrate [64]. The observed reduction of both zinc and citrate, is a clear indicator for loss of this characteristic prostate gland function in NCG spots with high CEG signature activity. This agrees with our ST-based findings of reduced transcriptional activity of citrate secretion associated genes in the same spots. This metabolic change of benign glands adjacent to cancer areas has, to our knowledge, not been previously observed in PCa. The loss of gene expression associated with citrate secretion in NCG spots from relapse cancer samples (**Figure 2e**) appears to be the most profound change of the prostate specific luminal cell-type characteristics. When inspecting the genes present in the RA signature, we found that two of the downregulated RA signature genes (*ALOX15B* and *NCAPD3*) were also central genes in our previously published network of genes for citrate secretion [44], further strengthening the link between aggressive tumors and loss of citrate secretion in corresponding cancer adjacent benign glands. Further, using independent publicly available data, we could support the associations between both signatures and reduced luminal functions, specifically citrate secretion on the transcriptional level. The observed increased glucose levels in the spatial data in combination with the increase lactate levels in bulk data in signature high scoring spots and samples are indicators for anaerobic glycolysis also known as the Warburg effect [63]. Further, build-up of AMP and reduction of ATP, resulting in an elevated AMP:ATP ratio, are indicative for a low energy state of the tissue and suggest high activity of anabolic processes [65]. This was accompanied by a potential shift to utilization of glutamate as an energy source as an adaptation to a high energy demand under hypoxic conditions suggested by the pronounced increase of NAA, glutamate, and glutamine [63]. All these metabolic changes observed in regions with high RA and CEG activity are typically observed in cancer or inflammation [22, 66]. The increased urate levels in these regions, indicative of a potential increased activity of the uric acid cycle, might be a further indicator for a high energy demand. Interestingly, urate, and urate crystals, have been implicated in being a cause for prostatitis [67-69] and a factor leading to inflammation eventually contributing to prostate carcinogenesis [70]. The increase of taurine in these areas is interesting as it has been described as pro-apoptotic, anti-prolific, and anti-migratory [71] but also to mitigate the resulting oxidative stress from inflammation, typically increased in high catabolic tissue [72, 73].

The bulk metabolomic data uncovered a comparable metabolic phenotype linked to increased signature activity as we found in and discussed for the spatially resolved data. Both signatures are associated with loss of normal prostate glandular function indicated by a reduction of citrate and indicators for an increased catabolism. Bulk results in the context of the very localized CEG active NCG must be interpreted carefully. Nevertheless, the results from bulk metabolite analysis are in support of the spatially resolved data obtained using MSI for both signatures and in agreement with previous work as we discussed in the previous paragraph. Of note, the increase of lactate, a metabolite we could only detect using bulk HRMAS NMR, in samples with high CEG activity, might suggest that increased anaerobic glycolysis in CEG active NCG regions could lead to accumulation of lactate. High intracellular levels of lactate have recently been reported to result in chromatin remodeling and to negatively affect the response to anti-androgen treatment in prostate organoids [74].

The results of this work suggest that early molecular changes happen in benign appearing glands of relapse patients displaying high RA and CEG activity resulting in loss of luminal characteristics such as citrate secretion, and an aberrant metabolism typical for PCa. These novel results, mapping very localized features of the PCa TME, support previous findings that the switch to a molecular cancer phenotype happens early in PCa before morphological changes typical for cancer are unambiguously identifiable in the tissue [8, 75, 76].

The RA signature developed from spatial analysis was validated by using bulk transcriptomics data, while the CEG signature lost its predictive power. Using ST data, we detected RA signature activity in all the major histopathology spot classes, including stroma and high-grade cancer, while the CEG signature was mainly active in some of the NCG regions in the TME. Thus, it appears reasonable that the predictive power of the RA signature is, to some extent, independent of the spatial information while the CEG signature requires precisely linked tissue morphology to unfold its full potential and should rather be perceived as a spatial, PCa-specific, TME marker. Using publicly available data, we found further indications that bulk transcriptomics data is limited compared to ST data in analyzing very localized gene expression-gen set associations. This clearly highlights the value of methods, like ST and MSI, with high-plexity and high spatial resolution to investigate otherwise hidden changes in the TME, while also making it clear that spatial technologies are required to fully explore the link between PCa relapse and these aberrant NCG regions. Simultaneously, this suggests that the number of patients and samples that were used to generate the spatial multi-omics data are a limitation specifically regarding generalizability of the spatial CEG signature for PCa relapse prediction. Nevertheless, the results provide novel insights and convincing evidence to justify future research on larger cohorts using our spatial multi-omics approach. This of course limits the immediate use of these signatures in the clinic if bulk samples were to be used for clinical testing. However, if the RNA-molecules or their protein products could be detected through multiplexed staining methods on tissue sections, these signatures hold potential to predict aggressive prostate cancer. Stained tissue sections are already routinely used in the clinic and capture the spatial context of heterogeneous PCa and future work should investigate the robustness of this strategy for clinical implementation.

### Conclusion

In this study, using spatial multi-omics data obtained from the same samples, we found two novel gene expression signatures capturing changes in the TME of PCa. We uncovered a strong link between Club-like cells, our signatures, and transcriptional and metabolic changes, specifically changes of citrate secretion in NCGs of patients with aggressive PCa. Further, we found indications that chemokines, major components in our signatures, are released by Club-like cells and are potentially responsible for the increased immune cell infiltration in and around these regions. In conjunction with the loss of citrate levels and a metabolism shifted towards higher energy consumption, these findings suggest a critical role of chemokines in the development and progression of PCa by modulating the TME. The increased expression of chemokines in PCa tissues highlights their potential as diagnostic and prognostic markers. Moreover, targeting chemokines and their receptors represents an attractive therapeutic strategy for combating PCa. Further research is warranted to unravel the intricate mechanisms underlying chemokine-mediated tumor progression, their role in altering prostate metabolism, and to explore their therapeutic potential fully.

## Methods

### Patient inclusion and sample collection

The study utilized human prostate tissue samples obtained from PCa patients who gave informed written consent before undergoing radical prostatectomy at St. Olav’s Hospital in Trondheim between 2008 and 2016. None of the patients had received any treatment prior to their surgery (further clinical details listed in Supplementary Table 1). This research received approval from the regional ethical committee of Central Norway (identifier 2017/576) and adhered to both national and EU ethical regulations.

In total, tissue samples from 37 patients were included in this study. Among them, ten patients remained relapse-free for >10 years following surgery, while twenty-seven patients experienced relapse (biochemical recurrence (PSA > 0.1 ng/ml) and/or confirmed metastasis). We collected 174 samples (no relapse n=48, relapse n=126) from all these 37 patients (no relapse n=10, relapse n=27). Of these samples, we selected for spatially resolved methods 32 samples (no relapse n=12, relapse n=20) from 8 of the 37 patients (no relapse n=3, relapse n=5) choosing relapse patients with confirmed metastasis within three years. All samples were obtained by taking a 2 mm thick slice from the middle of the prostate (transverse plane) immediately after surgical removal. This slice was snap frozen and stored at -80 °C following the procedure described by Bertilsson et al. [77]. The collection of tissue slices and their storage were conducted by skilled personnel at Biobank1® (St. Olav’s University Hospital in Trondheim). From each prostate slice we collected 8 to 13 tissue samples (3 mm in diameter) by using our in-house developed drilling device based on a previously described method [77]. For spatially resolved methods, we selected four samples per patient based on histopathology evaluation of hematoxylin-erythrosine-saffron (HES) stained tissue sections of each drilled sample. If applicable, we acquired two samples with cancerous tissue, one sample with non-cancerous morphology close to the cancerous region, and one sample with non-cancerous morphology distant from the cancerous area for each patient.

### Tissue serial sectioning

For all tissue samples we collected several 10 µm-thick sections through serial cryosectioning at -20°C using a Cryostar NX70 (Thermo Fisher). The tissue sections destined for spatial transcriptomics analysis were placed on Visium slides according to manufacturer’s recommendations. Remaining tissue sections were collected for other methodologies (many of which not presented in this study) and were placed on other types of slides such as super frost, conductive slides and membrane slides. Sections destined for matrix-assisted laser desorption ionization (MALDI) mass spectrometry imaging (MSI) were placed on conductive IntelliSlides (Bruker Daltonics) and vacuum-packed as described by Andersen et al. [78]. All sections were kept cold during sectioning (-20°C) and stored at -80°C until further use. Material remaining after sectioning was stored at -80°C until use for bulk metabolomics (HRMAS NMR) and transcriptomics (RNA-sequencing).

### RNA isolation, cDNA library generation and sequencing

For spatial transcriptomics profiling we used the Visium Spatial Gene Expression Slide & Reagent kit (10X Genomics) following the manufacturer’s manual. Shortly before RNA extraction, tissue sections were fixated with 100% methanol, stained with hematoxylin and eosin (HE) and digitally scanned at 20x magnification (Olympus VS120-S5). To ensure proper optical focus, a coverslip was temporarily placed on the tissue section and then gently removed after scanning. RNA was extracted by exposing the tissue sections to a permeabilization agent for 12 minutes. Optimal extraction time was previously determined with the Visium Spatial Tissue Optimization Slide & Reagent kit (10X Genomics). A second strand mix was added to create a complementary strand, and subsequently, cDNA was amplified using real-time qPCR. The amplified cDNA library was quantified using the QuantStudio™ 5 Real-Time PCR System (Thermo Fisher) through qPCR, and the cDNA libraries were stored at -20°C until further use. Paired-end sequencing was conducted using the Illumina NextSeq 500 instrument (Illumina®, San Diego, USA) and the NextSeq 500/550 High Output kit v2.5 (150 cycles). Bulk RNA-sequencing samples used in this study was performed as previously described [79].

### Generation of raw count tables

ST sample data were processed using 10x genomics space ranger software package (version 1.0.0) according to manufacturer’s recommendations. BCL files were demultiplexed to FASTQ files using spaceranger mkfastq. Raw count tables were generated from these FASTQ files together with the HE stained images and the human reference transcriptome GRCh38 version 3.0.0 per sample using spaceranger count. For the bulk samples, raw base call (BCL) files generated by Illumina sequencer were processed as previously described [79].

### Cell detection and number of cells per ST spot calculation

For each sample, cells were detected using the build-in-cell-detection feature of QuPath (version 0.2.3)[80] utilizing HE staining segmentation of nuclei to generate a list of nuclei centroids. For each spot, the number of centroids inside the spot area were counted to obtain the cells per spot count values. Data export from QuPath, merging with spatial transcriptomics spots, and cell counting were done using in-house developed groovy and python scripts and a python package is available on github [https://github.com/sekro/qupath_scripts, https://github.com/sekro/spatial_transcriptomics_toolbox]

### Histopathology

Two experienced uropathologists (T.V. and Ø.S.) independently evaluated the HE-stained sections from all 32 spatial transcriptomics samples in QuPath (version 0.2.3). They annotated cancer areas according to the International Society for Urological Pathology (ISUP) Grade Group (GG) system[81] and aggregates of lymphocytes. Glands of uncertain cancer status were also annotated. We aimed to assign pure Gleason patterns to distinct cancer glands where possible. For example, well-defined separate areas of Gleason pattern 3 and 4 were annotated as IUSP GG 1 and 4, respectively, instead of one area of ISUP GG 2 or 3 (Gleason 3+4 or 4+3). For the downstream data analysis, a consensus pathology annotation was reached in agreement with both pathologists. Gleason scores were transformed into ISUP GGs and labeled with ISUP1-5 accordingly. Cancer areas with uncertain grading were annotated as ISUPX (indecisive between ISUP3 and ISUP5) and ISUPY (indecisive between ISUP1 and ISUP4). Other section annotations included tissue borders, lumen, glands, stroma, and stroma with higher levels of lymphocytes (referred to as ‘lymphocyte-enriched stroma’).

### Integration of Histopathology with Spatial Transcriptomics data

Digital histopathology annotations including annotations for tissue bounds, luminal spaces, tissue folds, and uncertain areas were exported as binary images from QuPath using an in-house developed groovy script [https://github.com/sekro/qupath_scripts/blob/main/export_for_st_toolbox.groovy]. Histopathology data were merged with spatial transcriptomics data using an in-house developed python package [https://github.com/sekro/spatial_transcriptomics_toolbox]. For each spot, the intersection with the binary maps for each annotation class were calculated as a percentage per ST spot. The area of each of these intersections was divided by the tissue containing area of the spatial transcriptomics spot. The resulting percentage values per annotation class per ST spot were accumulated in a table and used to assign each spot a histopathology class. First, spots were excluded if they contained less than 50% tissue, more than 50% folded or low staining quality tissue, or more than 80% luminal space. If a spot contained more than 55% stroma it was assigned the stroma or lymphocyte-enriched stroma class using the one with the higher value. Otherwise, the spot was assigned one of the ISUP-grades, perineural invasion (PNI), or lymphocyte classes if the respective area of one of these classes was above 50%. In a last step, spots that were not assigned stroma or any of the ISUP-grades, PNI, or lymphocyte classes and contained at least 30% non-cancer glands (NCG), were assigned the class NCG. Spots not fulfilling any of the mentioned criteria were excluded from the analysis.

### Spatial Transcriptomics data curation and normalization by number of cells per spot

Raw counts were normalized by dividing the counts by the number of cells for each spot (**Figure 1a, Supplementary Figure 2**). Normalized total counts per spot were calculated as the sum of all normalized counts and normalized number of detected genes per spot were obtained by counting genes with normalized counts > 0 for each spot. Spots with less than 100 normalized total counts or less than 40 normalized number of detected genes per spot were excluded.

Genes were only included if total raw count per gene over all spots was not zero and at least 10 counts for at least 10 spots were observed.

### Identification of RA and Chemokine Enriched Gland (CEG) signatures

The ST spots were grouped according to the histopathology classes and analyzed for differences between relapsing and non-relapsing patients utilizing spatial information (see also **Supplementary Figure 1b**). ST data raw UMI counts per cell values were decadic logarithm transformed prior scaling by 10000 to obtain human readable values 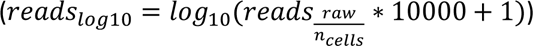 and used for all downstream analysis described in this section. To capture the variable genes in the ST data, we used two approaches: 1. We calculated for each histopathology class the relative standard deviation (RSD, standard deviation / sample mean) for each gene and saved the 5 genes with the highest RSD per class into a preliminary gene list. 2. To find genes that are variable compared to genes of similar average expression, we used the approach suggested by Satija et al. and calculated for each class the dispersion (mean/variance) for each gene [82]. Then, genes were binned according to their mean into 20 bins. Subsequently, dispersions were standardized by its respective genes bin mean and standard deviation to obtain bin-wise “expression-level normalized” z-scores for each dispersion. We added the 5 genes with the highest z-score dispersion per class to the preliminary gene list. Next, we added deregulated genes in relapsing patients per class to the preliminary gene list. Deregulated genes were defined as genes with an absolute difference of the means of the normalized expression of relapsing and non-relapsing patients larger than 1 (corresponding to a 10-fold change in expression) and a maximum RSD of 20% in the non-relapsing patient group.

The preliminary gene list was filtered by removing genes with 75% or more observations coming from only one sample or patient and that have been observed in 2.5% or less of all spots. Further, to rank the remaining genes according to their strength to differentiate between relapse and no relapse we calculated the min-max normalized zero-count-free median difference (ZFCMD) and the min-max normalized non-zero-counts spot count difference (NZCSCD), with

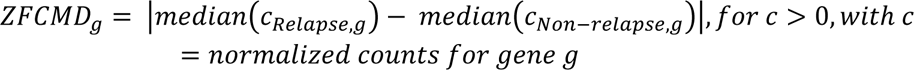

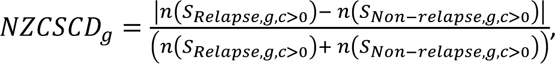 with S = spots, g = gene, c = normalized counts for gene g. By excluding spots with 0 counts the ZFCMD can better capture intensity differences in the sparse ST data but gets more sensitive to outliers in cases of low number of spots with counts above zero. Thus, we calculated the NZCSCD to obtain a measure of the frequency of spots with counts above zero and derived a gene rank by calculation the average of ZFCMD and NZCSCD.

Genes with a rank-score below the median of the observed rank-scores were removed from the preliminary gene list. The filtered gene list was analyzed for enriched gene ontologies (GO) using the GOATOOLS python package [83] showing enrichment of immune and chemokine related GOs (chemokine activity, chemokine-mediated signaling pathway, antimicrobial humoral immune response mediated by antimicrobial peptide, inflammatory response). Thus, we added all chemokines observed in the ST data to the preliminary list and re-ran the filtering to obtain the final filtered gene list. The RA signature was derived from this list by accumulating all genes that were unanimously up- or downregulated in all spot classes. The CEG signature was derived by accumulating genes that were exclusively up- or downregulated in relapse in NCG spots.

### Single-sample Gene Set Enrichment Analysis (ssGSEA) and score calculation

The ssGSEA approach is an in-house implementation of the algorithm presented by Markert et al. [24], which has also been used in a previous study by us on PCa samples[44]. To handle the sparsity challenge in ST data for ssGSEA, gene expression values were centered by subtracting the mean value for each gene before sorting. This will ensure that genes with zero expression in one sample will get a lower rank if it is highly expressed in other samples, compared to genes with zero or low expression across all samples. Scores were calculated separately for up- and downregulated parts of the RA and CEG signatures and combined by calculating the mean of the up- and negated downregulated ssGSEA scores. All ssGSEA scores were min-max scaled to a range between -1000 and 1000. Significant scores were determined by calculating a distribution of scores for n=1000 unique permutations of random gene rankings. One-sided p-values for each sample/spot score were calculated from this distribution and combined for up- and downregulated part using Fisher’s method. ST spot scores were considered significant when p-values <= 0.05/number of signature genes (p_RA_ <= 0.001, p_CEG_ <= 0.003) and an absolute score > 200. For bulk samples a p-values <= 0.05 and an absolute score > 200 was considered significant.

### Subclassification in 10 spot groups

Based on the classification of patients (relapse/no relapse), samples (cancer/field effect/normal) and spots (cancer, stroma, NCG) we assigned ST spots into 10 groups. We combined cancer and field effect samples into one cancer/field sample class and combined all ISUP-graded spots into one cancer spot class. Spots classified as (lymphocyte-enriched stroma, PNI, and others) were excluded from these spot-groups. Consequently, the 10 spot-groups were defined by having combinations of these classifiers: Level 1 (patient status): Relapse, no relapse (n=2), level 2 (sample class): Normal sample, cancer-field sample (n=2), level 3 (spot class): NCG, stroma (n=2 for level 2: Normal sample) or NCG, stroma, cancer (n=3 for level 2: Cancer-field sample)

### Deconvolution of spatial transcriptomics data

Each spot of the spatial transcriptomics data was deconvolved into estimated cell type fractions using stereoscope [25]. For single cell reference data we used publicly available human prostate single-cell RNA-sequencing data [10]. From this single-cell data we obtained a list of 4324 highly variable genes using scanpy’s implementation of the Seurat method [82, 84]. Stereoscope was run with this list of highly variable genes and the following parameters: sc epochs = 75000, sc batch size = 100, st epochs = 75000, st batch size = 100, learning rate = 0.01.

### Receptor-Ligand Interaction Analysis of Chemokines of RA and CEG signature

To investigate what receptors are present and might act as receivers of the chemokines observed in the RA and CEG sigantures squidpy’s v1.2.3 [85] implementation of the CellPhoneDB algorithm [86] was applied to the normalized and scaled but non-logarithm transformed (see section Identification of RA and Chemokine Enriched Gland (CEG) signatures) ST data. Omnipath [87] interactions were limited to “receptor” for the receiver and “ligand” transmitter and filtered to include only entries with at least one published reference. The histopathology classes assigned to the ST spots were used as cluster-key and permutation test was run with 1000 permutations and threshold of 0.01 for required cells per cluster. Values were calculated for each sample individually and corresponding interactions were subsequently combined by calculating the mean of the mean-values or combining the p-values using Fisher’s method. Not-a-number (NaN) p-values were omitted during combination. In case all p-values for one interaction were NaN, we set the respective p-value in the combined-result tables to 1. Results were visualized with squipy’s plotting function.

### Immunohistochemistry staining of LPS and LTA

To estimate bacterial infection/presence in the 32 ST samples we performed immunohistochemistry staining of neighboring 10 µm cryo-sections of lipopolysaccharides (LPS) and lipoteichoic acid (LTA). Cryo-sectioned tissue was fixed in 4% formalin for 15 min, washed twice in TBS-T at room temperature. Subsequently, peroxidases were blocked using 0.3% hydrogen peroxide followed by blocking of unspecific binding sites using in 10% normal serum with 1% BSA in TBS for 2 hours at room temperature. Primary antibodies against LPS and LTA (Abcam ab35654, Thermo MA1-7402) were used at 2,5 µg/ml and 2 µg/ml in TBS-T with 1 % BSA and incubated on tissue over night at 4°C. The tissue was washed twice in TBS-T prior incubation with secondary antibody (EnVision anti-mouse-HRP/DAB+ system, Agilent), 3,3′-Diaminobenzidine (DAB) development, counterstaining with hematoxylin, dehydration, and mounting were done according to manufacturer’s recommendations. Stained tissues were scanned on an Olympus VS200 ASW 3.3 (Build 24382) slide scanner in bright field at 40x magnification resulting in a resolution of 136,866 nm/pixel. Images were processed with QuPath using building functions for cell detection to obtain mean DAB stain intensity per cell. HE images of the ST data were registered onto corresponding DAB-stained images using affine transformation derived from 3 manually determined corresponding points for each image pair. For both RA and CEG signature, we calculated inverse-distance interpolated activity values for each cell center using the transformed ST-spot coordinates and signature activity as input (using k-dimensional tree, KDTree from python package SciPy, leafsize=10, use 8 nearest data points, eps=0.1)

### Metabolite detection with MALDI-MSI

Serial tissue sections from the same 32 samples that were used for ST were placed on conductive slides and analyzed with MALDI time-of-flight (TOF) MSI for the spatial detection of small metabolites and lipids. Prior to MSI, the sections were sprayed with matrix using the HTX TM-Sprayer^TM^ system (HTX Technology). The matrix solution consisted of 7 mg/mL *N*-(1-naphthyl) ethylenediamine dihydrochloride (NEDC) in 70% methanol. Spraying parameters included 18 layers with 0.06 ml/min flow rate and 1200 mm/min nozzle velocity, 3 mm track spacing, 10 psi pressure, 2 l/min gas flow rate, 40 mm nozzle height, 75°C nozzle spray temperature and 35°C plate temperature. Following matrix application, MSI was performed with a rapifleX™ MALDI TissuetyperTM (Bruker Daltonics) equipped with a 10 kHz laser shooting 200 shots per pixel at a 10 kHz frequency. Negative ion mode was used with a mass range of m/z 80-1000 and a spatial resolution of 30 µm. After MSI, the matrix-covered tissue sections were stored dry and dark at room temperature until HES staining was performed within 2 weeks. Metabolites were identified by MS/MS and accurate masses from our previous study using the same methods for equivalent PCa samples [88]. Zinc was detected as an adduct with chloride (ZnCl_3-_) and was previously identified by its isotopic pattern [21].

### Integration of ST and MALDI-MSI data

MALDI-MSI data were binned/down sampled to 80% of its original datapoints in FlexImaging (Version 5.0, Bruker Daltonics) and further baseline corrected (top-hat) and root-mean square (RMS) normalized using SCILS lab Pro (Version 2024a, Bruker Daltonics). Ion images of identified metabolites (intensity given by max peak height) were exported to imzML files and co-registered to ST data using our Multi-Omics Imaging Integration Toolset (MIIT) [27].

### HRMAS NMR Metabolomics

NOESY HRMAS NMR data was collected as previously described on a 600 MHz Bruker Avance III NMR spectrometer equipped with a magic angle spinning (MAS), ^1^H / ^13^C probehead (Bruker Biospin) [20]. After automics phase [89] and baseline correction, spectra were probabilistic quotient normalized [90], aligned with icoshift [91] and ppm ranges specific for the respective metabolites (succinate: 2.415-2.395, lactate: 4.15-4.08, citrate: 2.74-2.47, glutamate: 2.36-2.305, taurine: 3.445-3.37) numerically integrated using the trapezoidal rule. For each metabolite the raw integrals were min-max scaled.

### Validation of prognostic power of RA and CEG signatures in public data and comparison with prostate-related gene sets

We used 41 prostate-related gene sets selected from various sources [9, 10, 23, 44] in addition to our RA and CEG signatures for the ssGSEA analysis to evaluate prognostic power and relation to biological processes in the prostate of RA and CEG signatures (details in Supplementary Table S2). We used 38 of these gene sets for analysis in our ST and public data, and three additional gene sets for analysis only in public data (Supplementary Table S2 and S3). The three additional gene sets were made by genes for three commercially available gene signatures for aggressive PCa, CCP, GPS and Decipher [92]. For these three, we used only the selection of genes from these signatures and not the scoring algorithms for scoring the signatures in clinical samples.

Prostate specific-cell type and immune-cell genes downloaded from Joseph et al 2021 (Supplementary Table S3 and S8) [10] contained genes enriched in 5 prostate-specific cell types, and prostate-specific enrichment of genes in 19 immune-cell types. To created prostate-specific cell type gene sets from these 24 cell types, we ranked the genes for each cell type by q-value (ascending) and fold-change (descending). The genes were sorted by the average rank over q-value and fold-change, and the top 75 ranked genes were selected to create a gene set for each cell type. We further validated each of the 19 immune-cell type gene sets with 142 corresponding immune gene sets downloaded from Azimuth (https://azimuth.hubmapconsortium.org/) [93], for PBMC (Peripheral Blood Mononuclear Cells) version 1-3, and Bone Marrow version 1 and 2. Each gene set from Azimuth contained 10 genes. We used GSEA [24] to score each of the 164 Azimuth gene sets in the 19 ranked immune gene sets from Joseph et al. [10], and then evaluated whether gene sets from Azimuth were enriched in our gene sets generated from Jospeh et al. [10] The results generally showed enrichment of Azimuth immune gene sets with corresponding immune gene sets from Joseph et al., demonstrating that the gene sets downloaded from Joseph et al. represented their respective immune cell-types well. Two exceptions were CD4+ Naïve and CD8+ Naïve signatures which showed a more general enrichment in various CD4+ and CD8+ cells from Azimuth. The validation was also not able to distinguish subtypes of immune cells, for example different Dendritic subtypes (EREG, IGSF21, and TREM2), Basophil-Mast 1 and 2, Classical and Nonclassical Monocyte, and Natural Killer and Natural Killer T cells.

To compare these 41 gene sets and evaluate prognostic power and relation to biological processes in the prostate of our RA and CEG signatures in public data, we calculated ssGSEA scores for all gene sets in all 2512 samples from 12 public datasets (Supplementary Table S3). The public data gene expression values were not centered before ssGSEA analysis. Correlations between gene sets in each dataset were calculated by Pearson correlation. The overall correlation for two gene sets was calculated as the average correlation over all 12 datasets. Complete tables with correlations for all gene sets in each individual dataset are given in Supplementary File 3.

## Supporting information

Supplementary File 2 - GOE results

Supplementary File 3 - Public Data

## Data availability

Bulk and spatial transcriptomics data are available upon request through Federated European Genome Phenome Archive (FEGA) Norway data access committee with accession number EGAC50000000277 and bundled under study EGAS50000000413.

## Acknowledgement

This research was funded by the European Research Council (ERC) under the European Union’s Horizon 2020 research and innovation program (grant agreement no. 758306), Norwegian University of Science and Technology (NTNU), the Liaison Committee between the Central Norway Regional Health Authority (RHA) and NTNU, the Norwegian Cancer Society and Terje Eugen Johnsen funds. May-Britt Tessem’s research project is part of the Centre for Digital Life Norway, which is supported by the Research Council of Norway’s grant 248810. All tissue samples were collected and stored by Biobank1, St. Olav’s Hospital. Tissue sectioning, staining and scanning were performed by or in collaboration with the Histology lab at the Cellular & Molecular Imaging Core Facility (CMIC) at NTNU. The authors thank Ingunn Nervik, Kathrin Juanita Gravvold Torseth, and Borgny Ytterhus for excellent technical assistance during these experiments. Biobank1 performed isolation of RNA before bulk analysis. Transcriptomics experiments were carried out at the Genomics Core Facility at NTNU. MALDI MSI and HRMAS NMR acquisitions were achieved using instrumentation at the MR Core facility, NTNU. Alfonso Urbanucci wishes to thank the Academy of Finland project no. 349314; Cancer Foundation Finland; Norwegian Cancer Society project no. 198016-2018 & project no. 333723-2023, Tampere Institute for Advanced Study.

