## Supplementary Figure and Tables for "Deep phenotyping of the prostate tumor microenvironment reveals molecular stratifiers of relapse linked to inflammatory chemokine expression and aberrant metabolism"

#### Content list

Supplementary Figure 1: Histopathology classes and signature identification flow chart.

Supplementary Figure 2: Spatial Transcriptomics data normalization by number of cells per spot.

Supplementary Figure 3: Spatial distribution of relapse and NAG signature activity.

Supplementary Figure 4: Histopathology class composition of spots grouped by signature activity.

Supplementary Figure 5: Gene signature activity distribution in ST data.

Supplementary Figure 6: Spearman correlation of histopathology and cell types.

Supplementary Figure 7: Spearman correlation of all Relapse and NAG signature genes with cell types.

Supplementary Figure 8: Chemokine receptor expression detected in spatial and bulk transcriptomics data.

Supplementary Figure 9: Lipoteichoic acid (LTA, gram-positive bacteria) and lipopolysaccharides (LPS, gram-negative bacteria) staining results.

Supplementary Figure 10: Relapse signature score distribution in bulk samples from own cohort.

Supplementary Figure 11: NAG signature score distribution in bulk samples from own cohort.

Supplementary Table 1: Clinical data of all 37 patients with prostate tissue transcriptomics profiling in this study.

Supplementary Table 2: List of gene sets used for ssGSEA analysis.

Supplementary Table 3: Data sets used for analysis in public data.

### Supplementary Figures

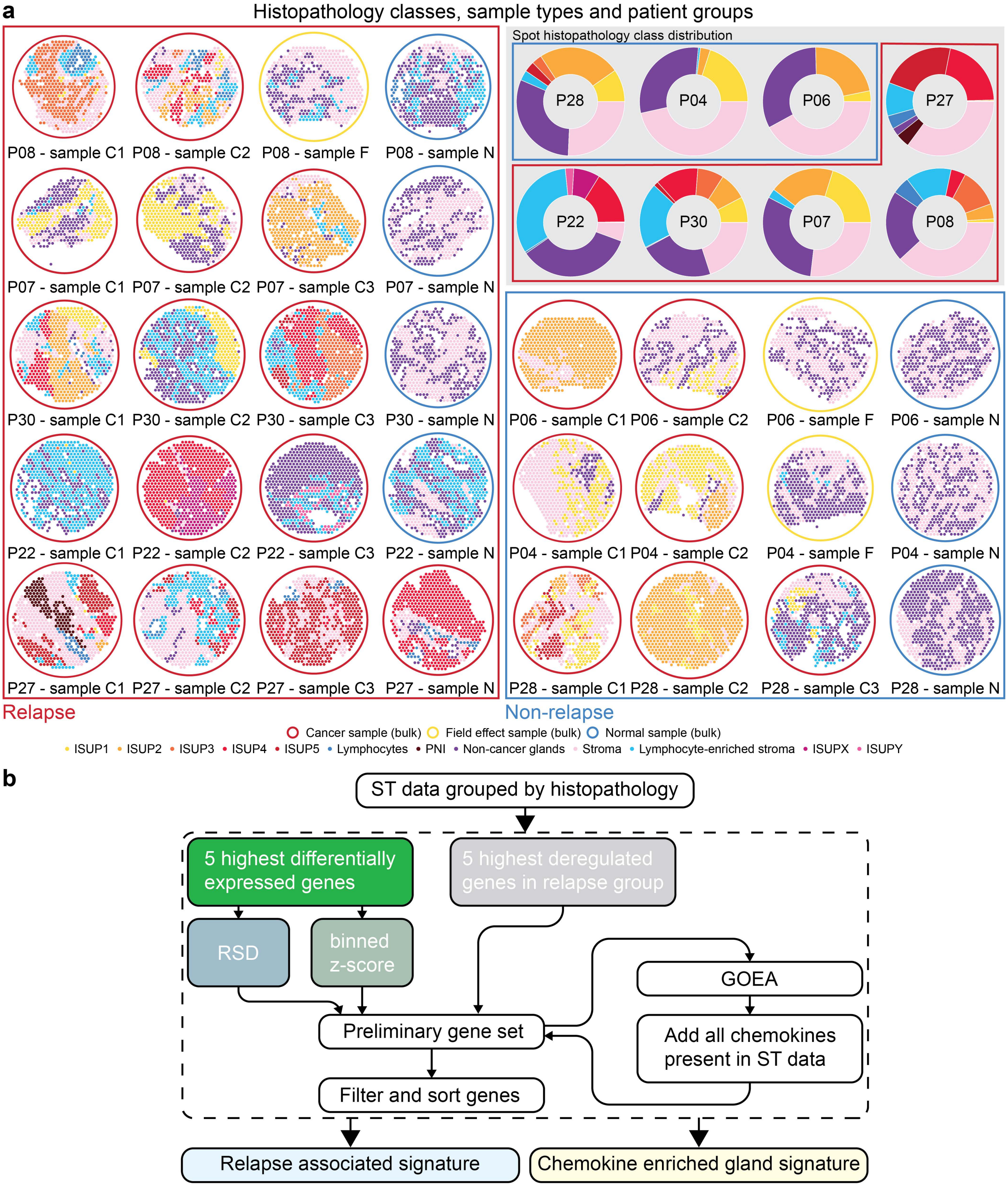


*Supplementary Figure 1 Histopathology classes and signature identification flow chart. (a) Histopathology classes visualized for all spots of all samples grouped by patient relapse status (left, red box = relapse, right, blue box = Non-relapse), sorted by patients (rows) and sample type (columns: C = cancer, F = field effect, N = normal). Insert top right is showing the histopathology class distribution for each patient grouped by relapse status (blue = non-relapse, red = relapse). (b) Flow chart illustrating the algorithm used to generate the two signatures: Relapse associated (RA) and Chemokine enriched gland (CEG).*

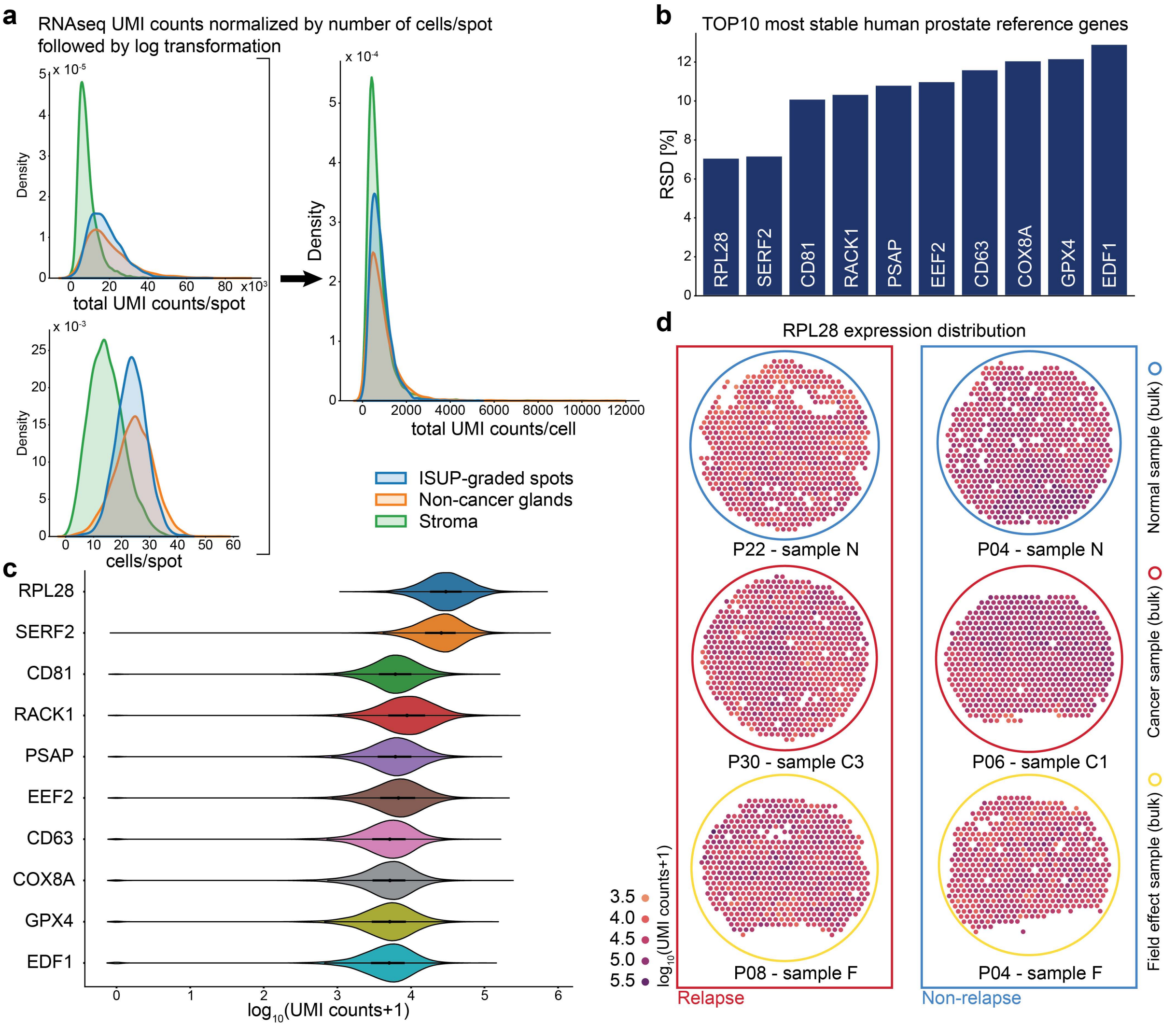


*Supplementary Figure 2 Spatial Transcriptomics data normalization by number of cells per spot. (a) The total UMI reads/spot and number of cells/spot were increased in cancer and NCG spots as compared to stroma spots. Scaling the reads by number of cells per spot results in spot-type independent total UMI reads distributions. (b) The relative standard deviation (RSD) of 10 select reference genes for the human prostate were all below 13% after normalization by cells/spot and showed comparable intensity (c) and spatial (d) distributions after log10 variance stabilization over all samples.*

**a** Relapse associated (RA) signature spatial distribution

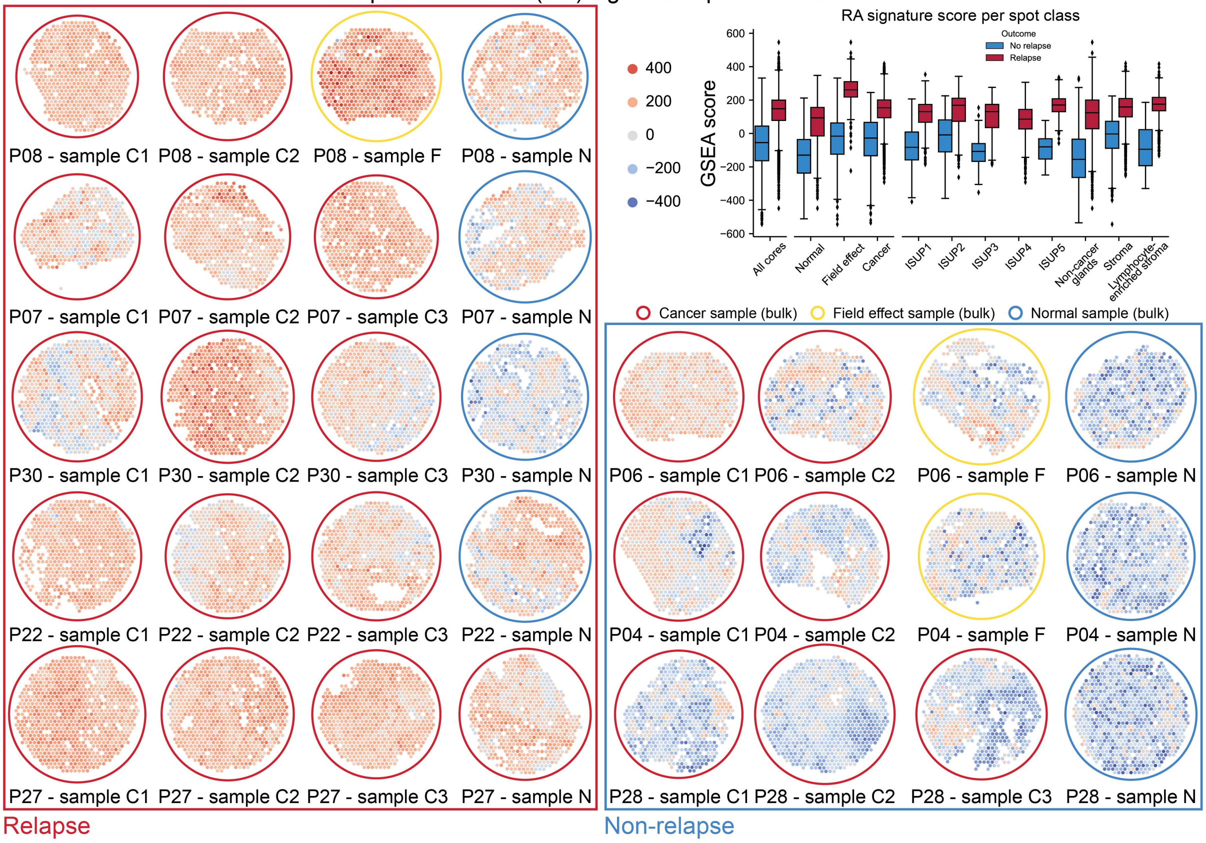


**b** Chemokine enriched gland (CEG) signature spatial distribution

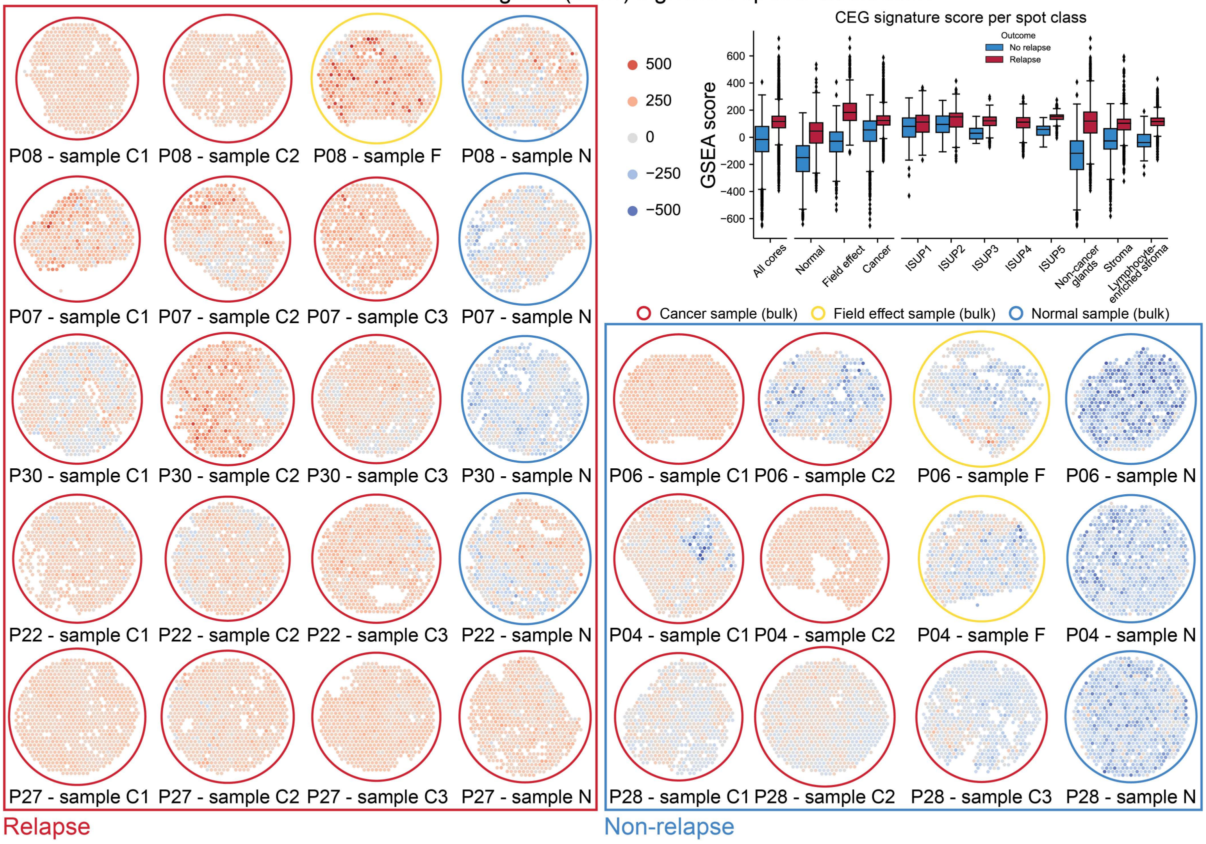


*Supplementary Figure 3 Spatial distribution of relapse associated (RA) and chemokine enriched gland (CEG) signature activity. (a) Distribution of the RA and (b) CEG score in all ST samples grouped by patient relapse status (left, red box = relapse, right, blue box = Non-relapse), sorted by patients (rows) and sample type (columns: cancer, field effect, normal). Inserts show*

*respective score distributions as box-and-whisker plots separated by relapse status (blue = no relapse, red = relapse) for all cores/samples and grouped by core type (normal, field effect, cancer), and histopathology classes (ISUP1-5, NCG, stroma, lymphocyte-enriched stroma)*

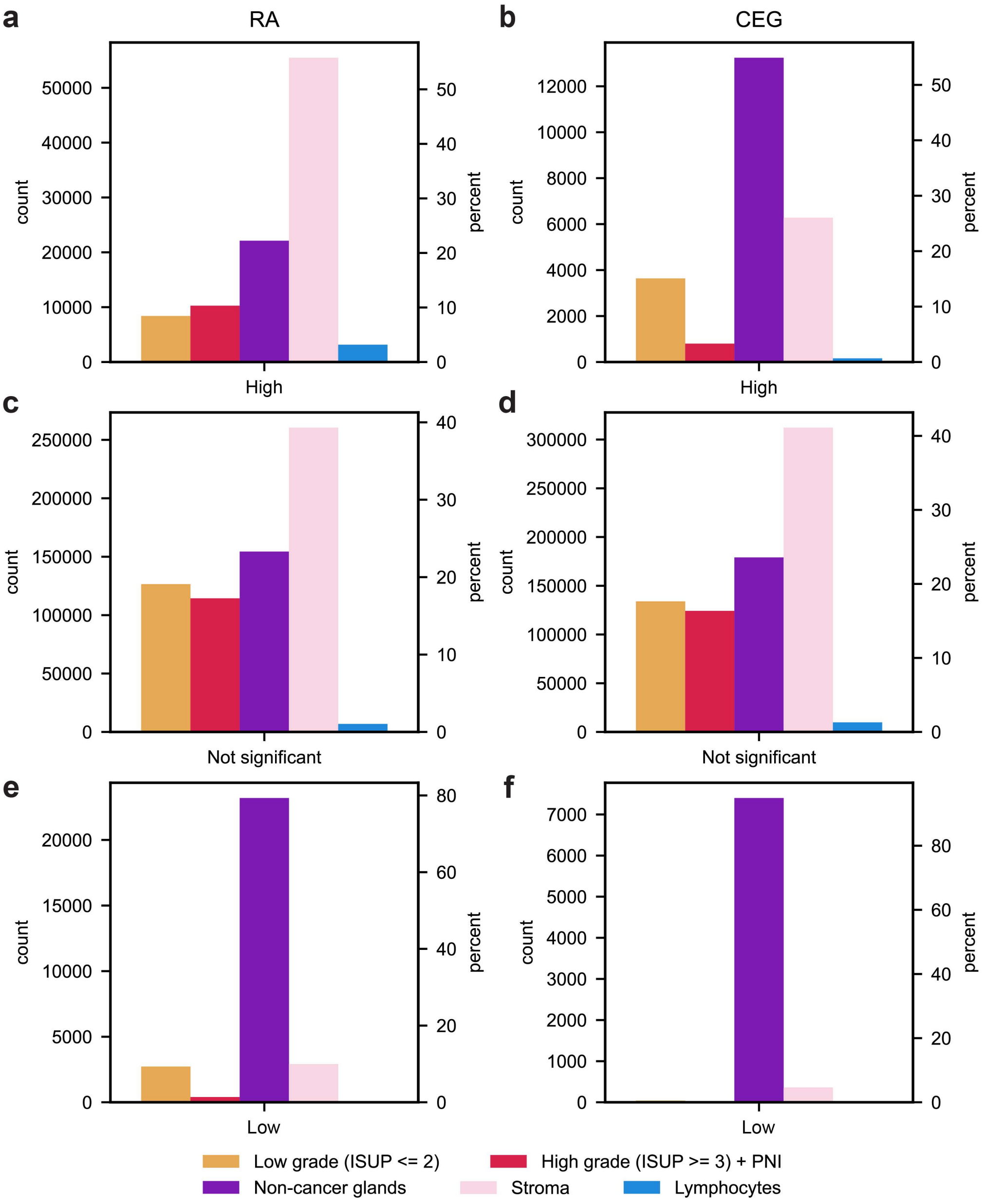


*Supplementary Figure 4 Histopathology class composition of spots grouped by signature activity. The compositions are shown for (a) high, (c) not significant, (d) low RA and (b) high, (d) not significant, (f) low CEG signature scoring spots as spot counts and percentage of all spots of the respective signature scoring group. Cancer spots were grouped into low grade (all spots classified as ISUP1-2) and high grade (all spots classified as ISUP3–5 and PNI).*

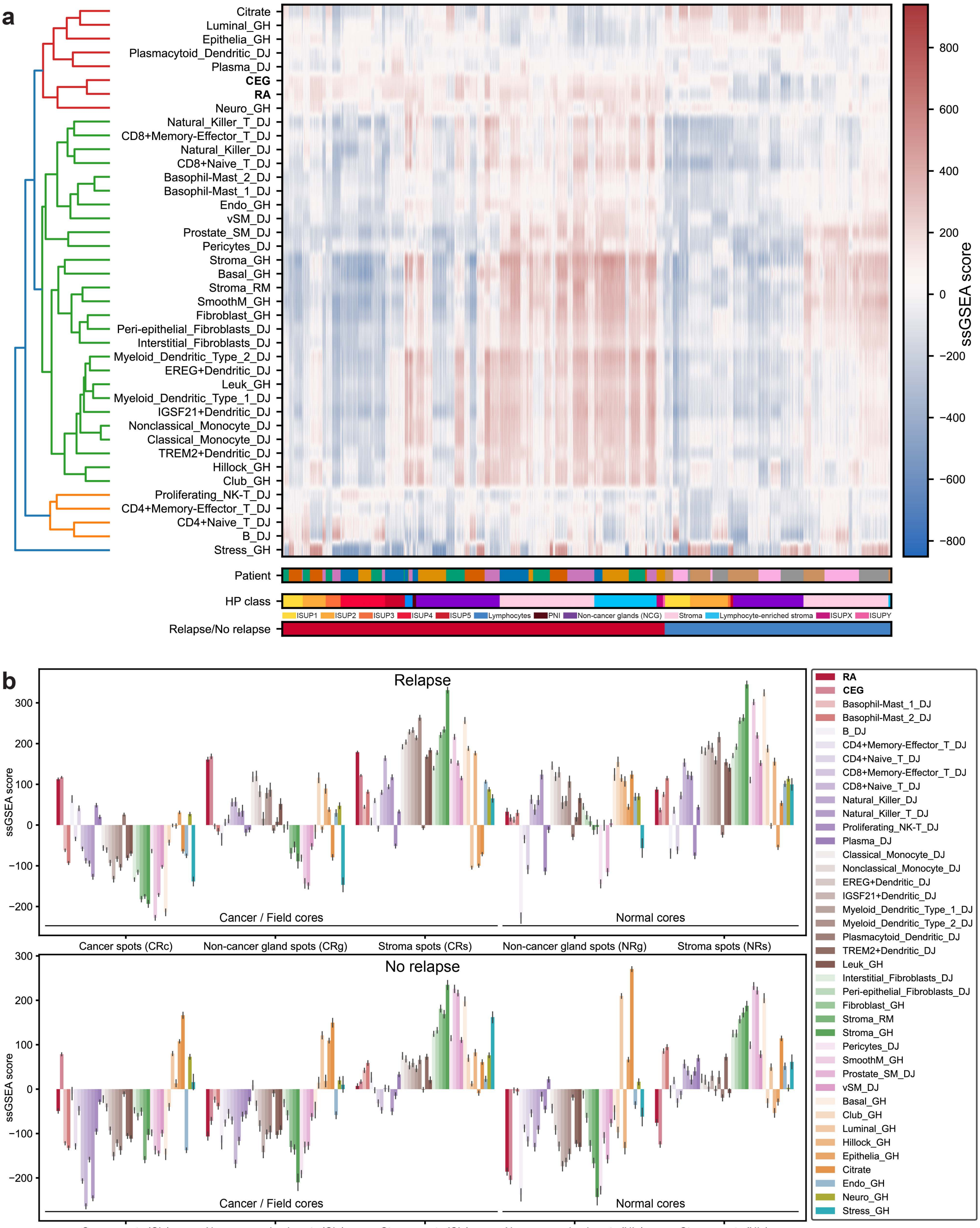


*Supplementary Figure 5 Gene signature activity distribution in ST data. (a) Signature scores are visualized for each spot by color, grouped as indicated by color bars in the 3 rows at the bottom by relapse status (red = relapse, blue = no relapse), histopathology (HP), and patients. Gene sets/signatures were clustered according to distribution similarity to enhance readability. (b) Mean (+/- standard deviation) activity of all prostate-related gene sets, cell type specific gene set, RA, and CEG signature of ST data spots group according to sample type (cancer/field or normal), relapse status, and spot class (Non-cancer gland [NCG], cancer, stroma).*

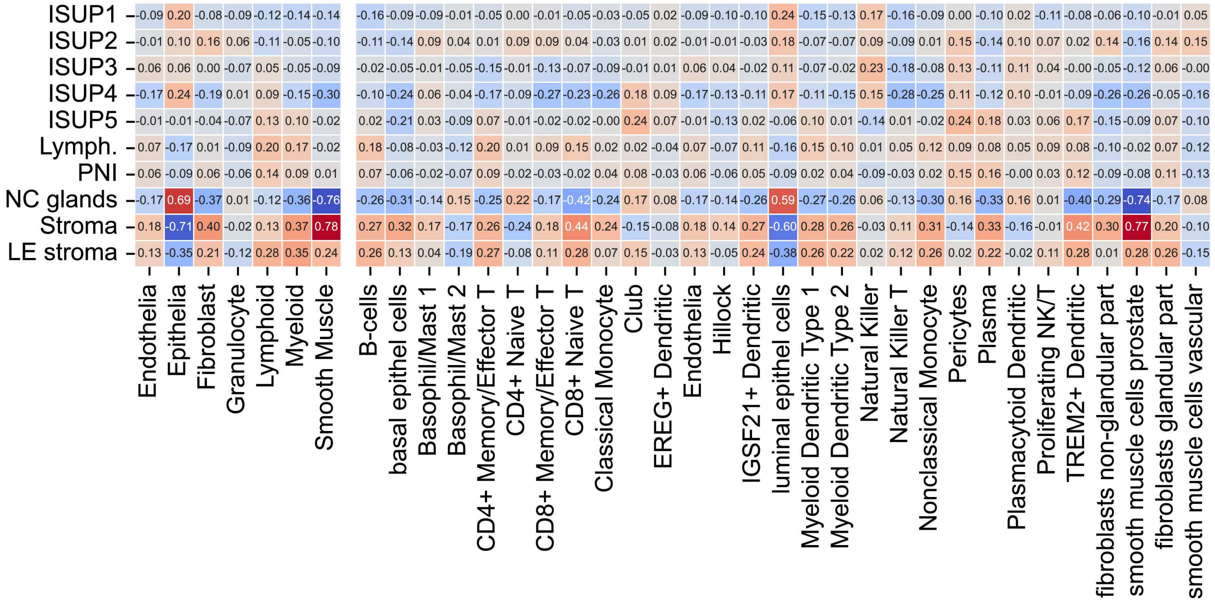


*Supplementary Figure 6 Spearman correlation of histopathology and cell types. Correlation coefficients are given as numbers and additionally visualized by color (blue = negative, red = positive correlation). Correlation coefficients were calculated using the raw histopathology class fractions per spot (see Methods) and cell type fractions obtained by spot deconvolution.*

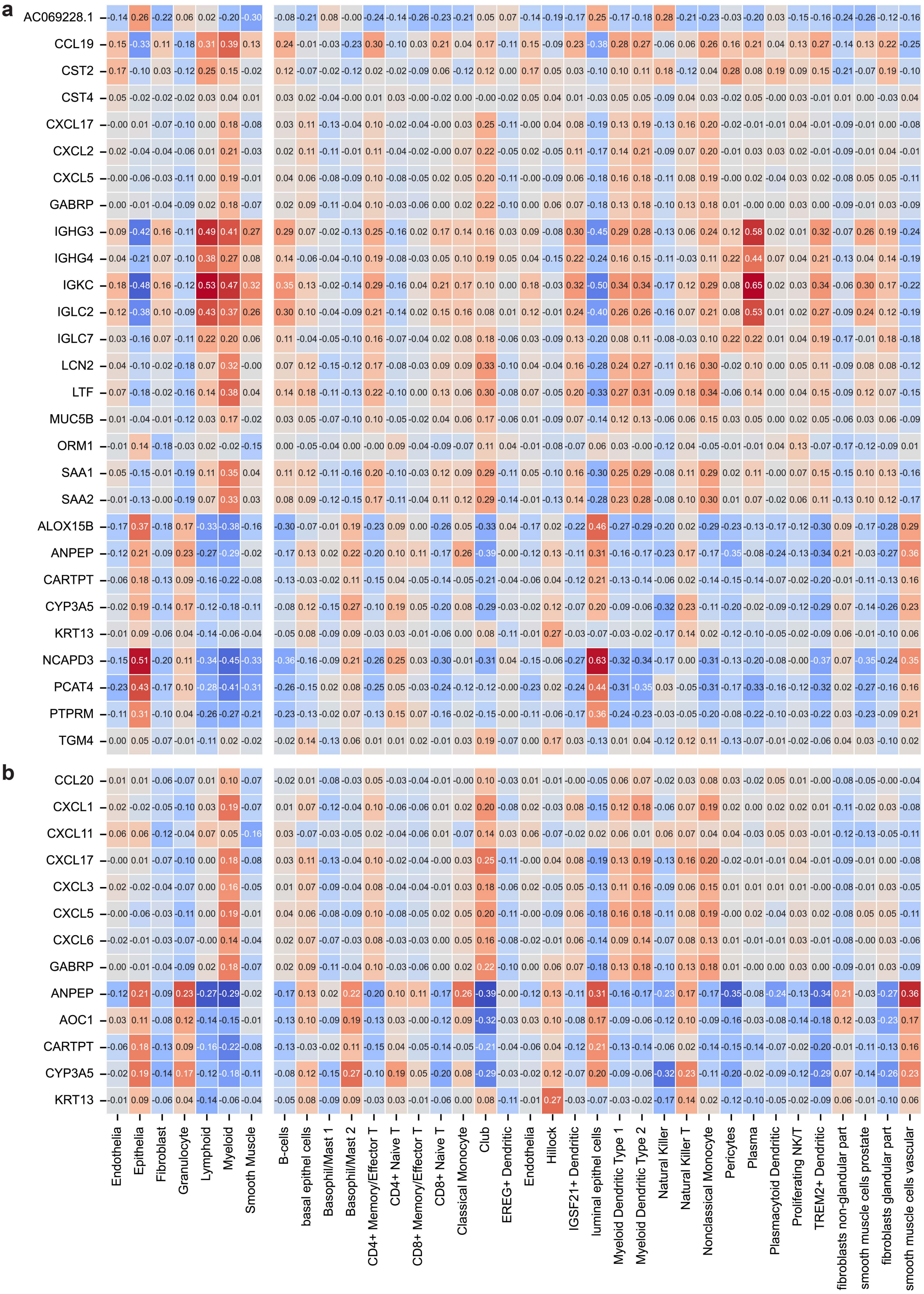


*Supplementary Figure 7 Spearman correlation of all RA (a) and CEG (b) signature genes with cell types. Correlation coefficients are given as numbers and additionally visualized by color (blue = negative, red = positive correlation). Correlation coefficients were calculated using the normalized reads per spot (see Methods) and cell type fractions obtained by spot deconvolution.*

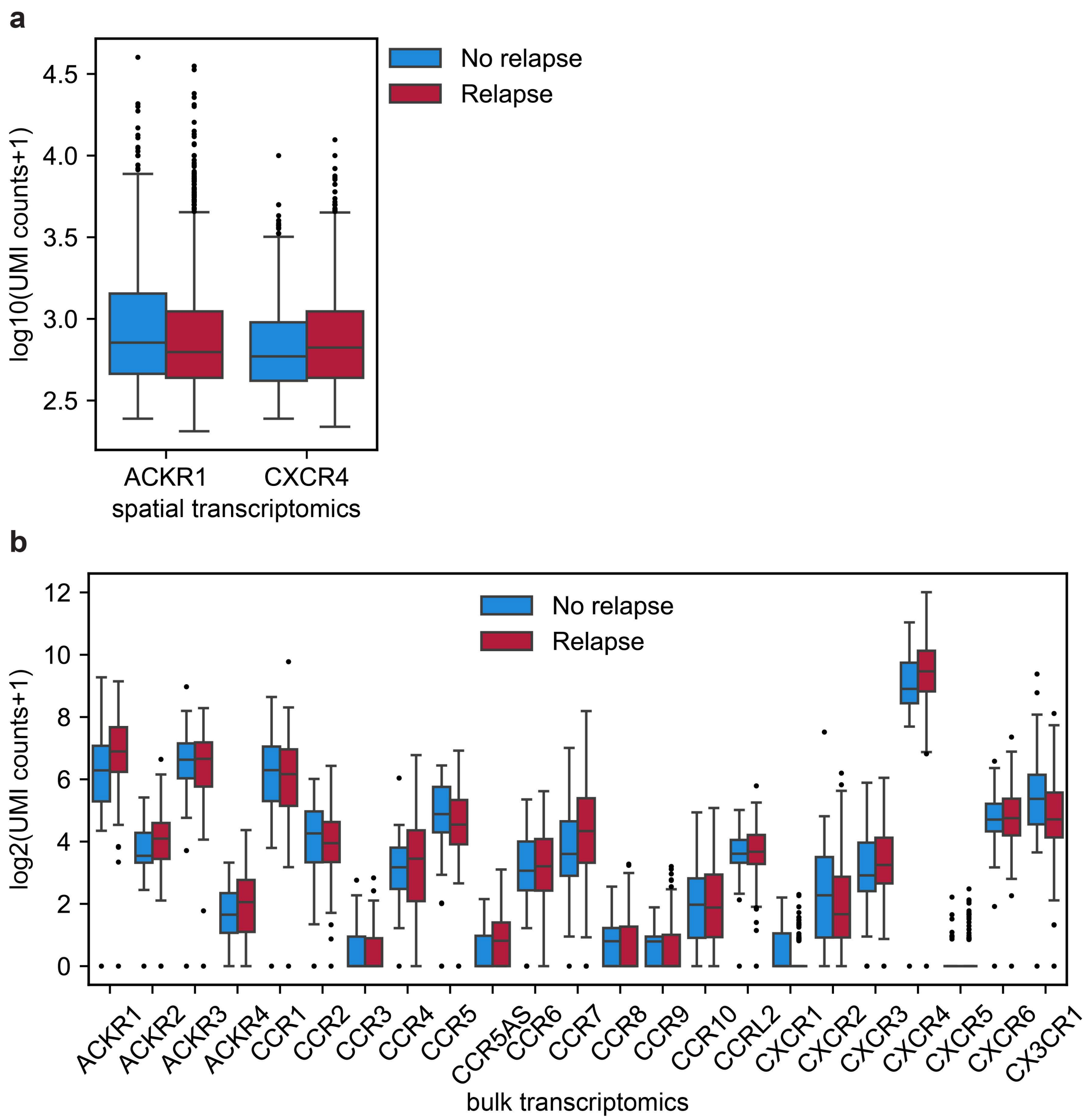


*Supplementary Figure 8 Chemokine receptor expression detected in spatial and bulk transcriptomics data. (a) Logarithm-transformed normalized expressions obtained by (a) spatial transcriptomics of 32 samples (no relapse n=12, relapse n=20) from 8 (no relapse n=3, relapse n=5) patients and (b) bulk transcriptomics of 174 samples (no relapse n=48, relapse n=126) from 37 patients (no relapse n=10, relapse n=37) are shown as box-and-whisker plots separated by relapse status (blue = no relapse, red = relapse).*

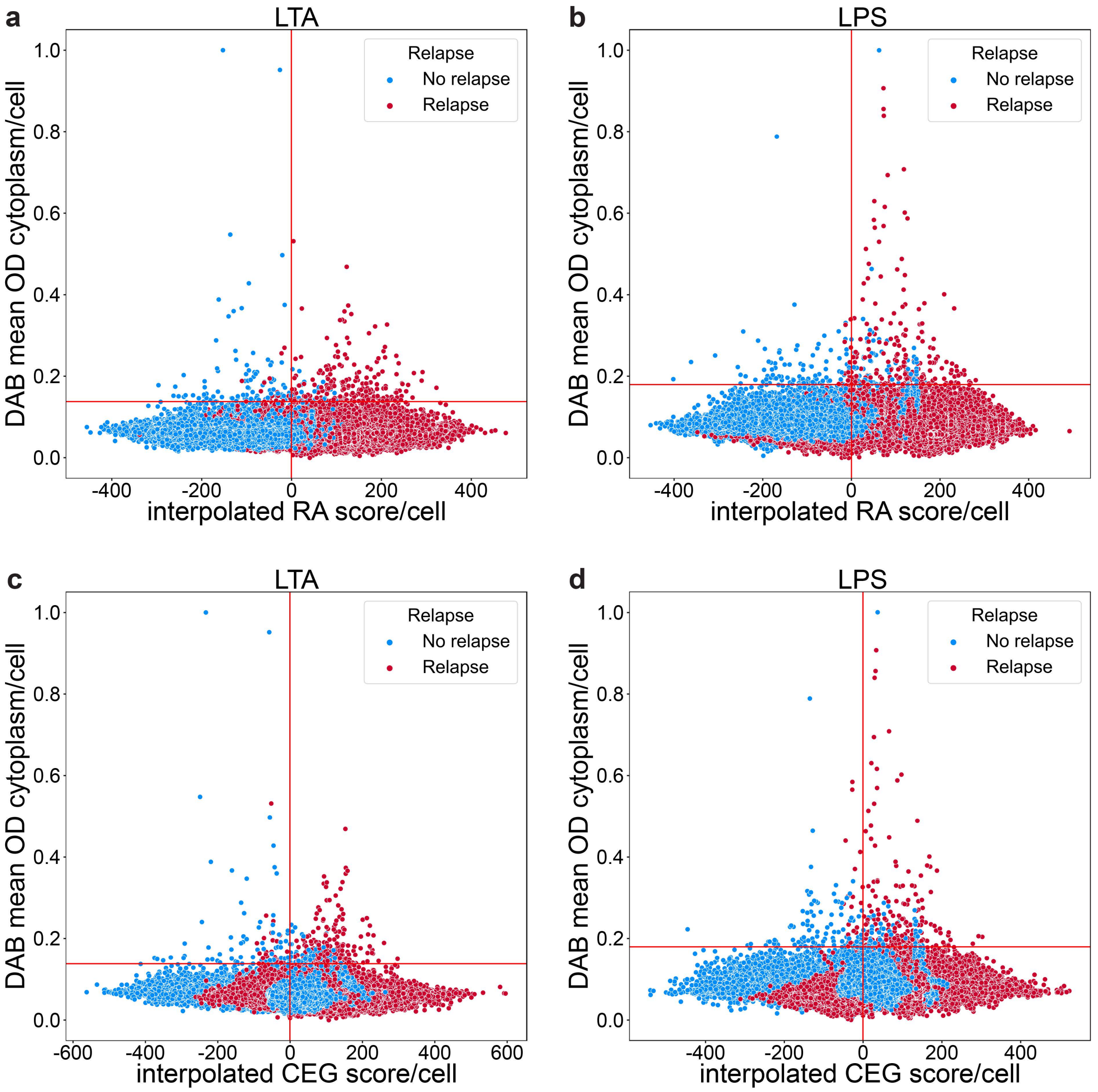


*Supplementary Figure 9 Lipoteichoic acid (LTA, gram-positive bacteria) and lipopolysaccharides (LPS, gram-negative bacteria) staining results. LTA (a, c) and LPS (b, d) cells computationally extracted from stained tissue were assigned mean DAB optical density (OD) values min-max normalized between 0 and 1 based on the staining intensity and an interpolated RA (a, b) and CEG (c, d) signature score (see Methods for details). Every cell detected is visualized as a point in the scatter plots grouped by relapse status (blue = no relapse, red = relapse). Horizontal red lines indicate positive DAB staining cutoff.*

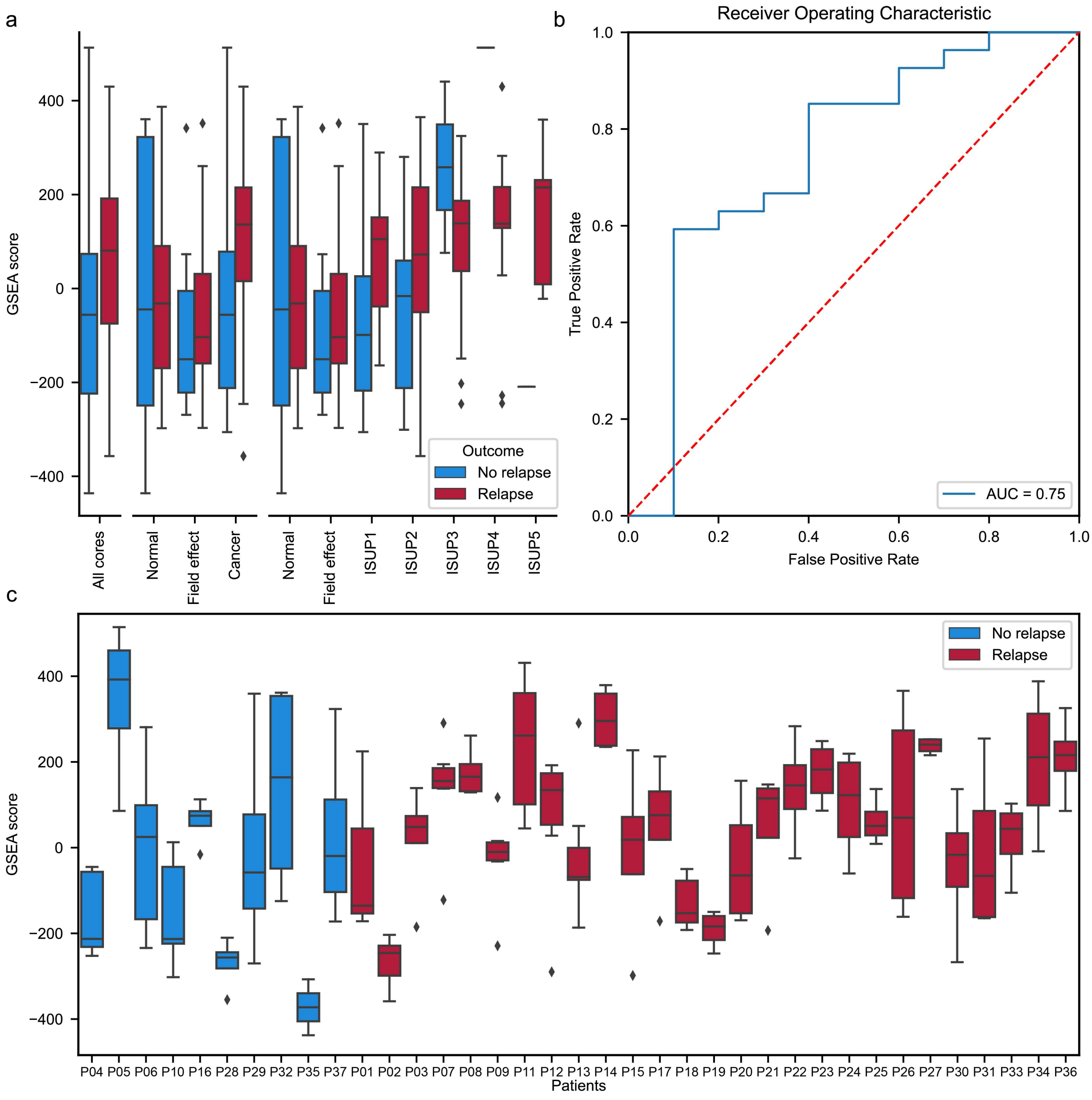


*Supplementary Figure 10 RA signature score distribution in bulk samples from own cohort. (a) Distributions are shown as box-and-whisker plots separated by relapse status (blue = no relapse, red = relapse) and shown for all cores, grouped by core/sample type (normal, field effect, cancer), and sample/core type and ISUP grade (normal, field effect, ISUP1-5). (b) Receiver operating characteristics (ROC) of the relapse signature in bulk samples resulted in an area under the curve (AUC) of 0.75. (c) Signature score distribution shown as box-and-whisker plots of all samples per patient grouped by relapse status (blue = no relapse, red = relapse).*

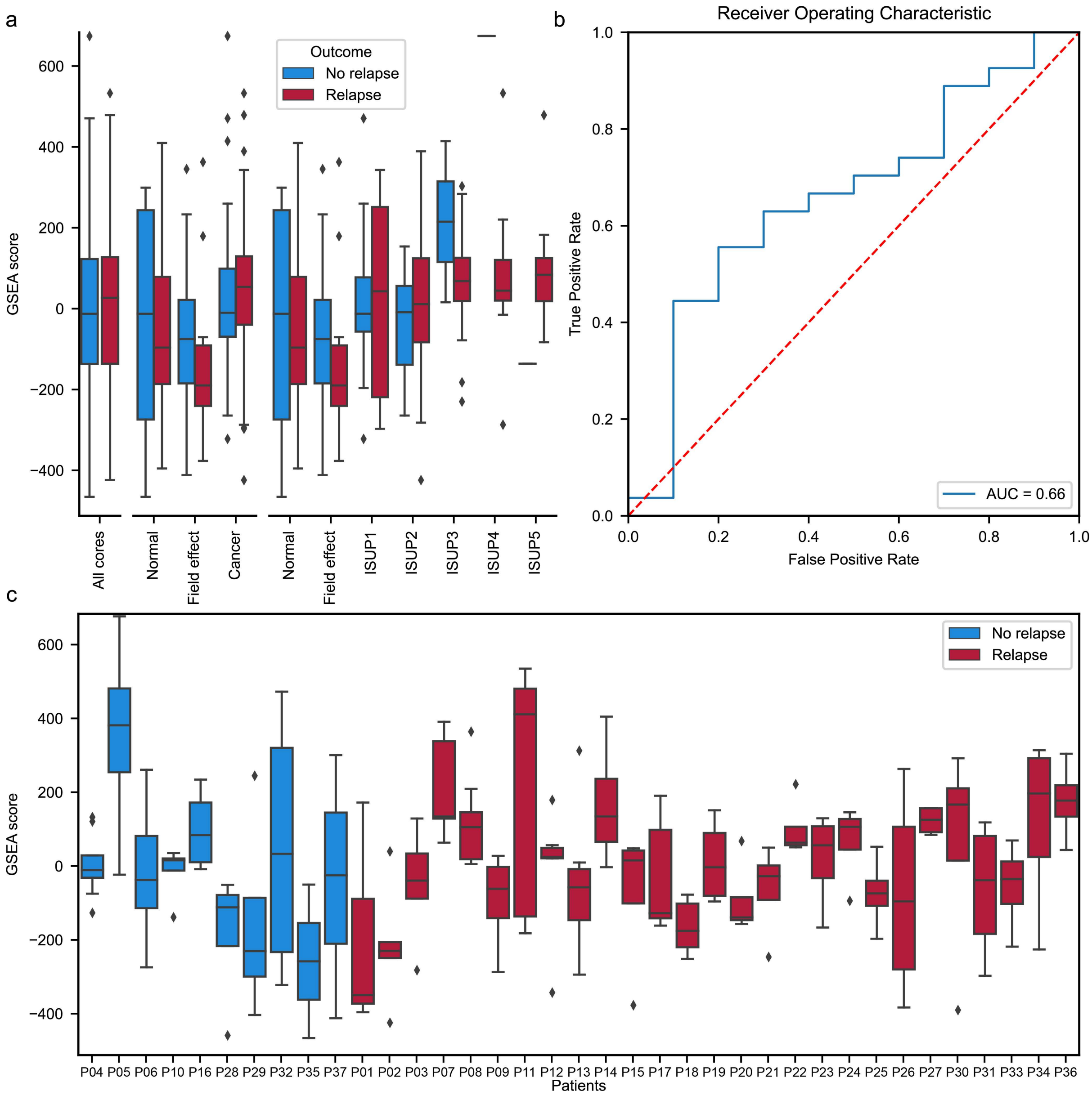


*Supplementary Figure 11 CEG signature score distribution in bulk samples from own cohort. (a) Distributions are shown as box-and-whisker plots separated by relapse status (blue = no relapse, red = relapse) and shown for all cores, grouped by core/sample type (normal, field effect, cancer), and sample/core type and ISUP grade (normal, field effect, ISUP1-5). (b) Receiver operating characteristics (ROC) of the NAG signature in bulk samples resulted in an area under the curve (AUC) of 0.66. (c) Signature score distribution shown as box-and-whisker plots of all samples per patient grouped by relapse status (blue = no relapse, red = relapse).*

### Supplementary Tables

*Supplementary Table 1 Clinical data of all 37 patients with prostate tissue transcriptomics profiling in this study. Patients with both spatial transcriptomics and bulk transcriptomics profiling are marked in bold.*

| ID | Status | Time to relapse (months) | Follow up time w. PSA | Total follow up time | Age at operation | Mean preoperative PSA | Preoperative Grade group | Clinical T-stage | EAU risk group for biochemical recurrence** | Operative Grade group | Post op. T-stage |
|---|---|---|---|---|---|---|---|---|---|---|---|
| P01 | Relapse | 32 | 78 | 78 | 62 | 7.3 | 5 | T3 | H | 4 | T3a |
| P02 | Relapse | 0 | 102 | 102 | 68 | 9 | 3 | T1c | I | 3 | T2c |
| P03 | Relapse | 0 | 62 | 62 | 58 | 47.9 | 3 | T3b | H | 5 | T3b |
| **P04** | **Control** | **135** | **12** | **135** | **63** | **10** | **2** | **T2** | **I** | **2** | **T2c** |
| P05 | Control | 122 | 9 | 122 | 66 | 5.4 | 2 | T2 | I | 2 | T2c |
| **P06** | **Control** | **136** | **15** | **136** | **56** | **9.6** | **2** | **T1c** | **I** | **2** | **NA*** |
| **P07** | **Relapse** | **37** | **107** | **107** | **59** | **12.8** | **3** | **T2** | **I** | **2** | **T3b** |
| **P08** | **Relapse** | **0** | **78** | **78** | **57** | **11** | **5** | **T2b** | **H** | **5** | **T3a** |
| P09 | Relapse | 0 | 62 | 62 | 51 | 26.4 | 5 | T3b | H | 5 | T3b |
| P10 | Control | 123 | 14 | 123 | 59 | 16.4 | 2 | T2/3 | H | 2 | T2c |
| P11 | Relapse | 14 | 79 | 79 | 57 | 18.4 | 2 | T2a | I | 3 | T3b |
| P12 | Relapse | 114 | 141 | 141 | 67 | 9.8 | 3 | T3a | H | 3 | T2c |
| P13 | Relapse | 11 | 56 | 56 | 65 | 19 | 1 | T1c | I | 2 | T2c |
| P14 | Relapse | 0 | 2 | 2 | 67 | 6.2 | 5 | T3b | H | 5 | T3b |
| P15 | Relapse | 1 | 33 | 33 | 59 | 60 | 5 | T3b | H | 4 | T3b |
| P16 | Control | 138 | 32 | 138 | 66 | 6.4 | 2 | T2 | I | 2 | T2c |
| P17 | Relapse | 19 | 70 | 70 | 63 | 9.5 | 2 | NA* | I | 2 | T3a |
| P18 | Relapse | 44 | 50 | 50 | 63 | 5.9 | 2 | T2c | H | 2 | T2c |
| P19 | Relapse | 0 | 93 | 93 | 65 | 4.6 | 4 | NA* | H | 2 | T2c |
| P20 | Relapse | 15 | 102 | 102 | 54 | 13.9 | 2 | T2 | I | 2 | T2c |
| P21 | Relapse | 0 | 39 | 39 | 66 | 11.4 | 5 | T3a | H | 5 | T3b |
| **P22** | **Relapse** | **0** | **34** | **34** | **53** | **32.5** | **3** | **T3a** | **H** | **4** | **T3a** |
| P23 | Relapse | 45 | 53 | 53 | 68 | 11.6 | 3 | T2c | H | 3 | T3a |
| P24 | Relapse | 25 | 66 | 66 | 65 | 4.8 | 3 | T2b | I | 3 | T3a |
| P25 | Relapse | 2 | 48 | 48 | 69 | 15.1 | 4 | T2b | H | 4 | T3b |
| P26 | Relapse | 107 | 125 | 125 | 60 | 7.7 | 2 | NA* | I | 3 | T2c |
| **P27** | **Relapse** | **0** | **44** | **44** | **73** | **16.3** | **5** | **T3b** | **H** | **5** | **T3b** |
| **P28** | **Control** | **119** | **6** | **119** | **63** | **22.3** | **2** | **T2** | **H** | **3** | **T2c** |
| P29 | Control | 133 | 11 | 133 | 55 | 10.2 | 2 | T3 | H | 2 | T2c |
| **P30** | **Relapse** | **0** | **22** | **22** | **53** | **45.9** | **4** | **T2c** | **H** | **3** | **T2c** |
| P31 | Relapse | 45 | 45 | 45 | 55 | 52.5 | 2 | T3 | H | 2 | T3b |
| P32 | Control | 122 | 20 | 122 | 59 | 11.5 | 2 | T1c | I | 2 | T2c |
| P33 | Relapse | 7 | 44 | 44 | 73 | 7.3 | 2 | T2b | I | 3 | T3b |
| P34 | Relapse | 38 | 83 | 83 | 66 | 8.4 | 2 | T2c/T3 | H | 3 | T2c |
| P35 | Control | 128 | 11 | 128 | 56 | 5.6 | 2 | T1c | I | 2 | T2c |
| P36 | Relapse | 0 | 30 | 30 | 70 | 29 | 3 | T3a | H | 3 | T3b |
| P37 | Control | 140 | 2 | 140 | 51 | 10.2 | 5 | T3 | H | 3 | T2c |

** Information was not available.*

*** EAU intermediate = I, high = H risk group for biochemical recurrence according to EAU based on mean preoperative.*

*Supplementary Table 2: List of gene sets used for ssGSEA analysis. References below this Table.*

| Gene set name | Number of genes | Genes in ST data | Source | Comment |
|---|---|---|---|---|
| B_DJ | 75 | 54 | Joseph, 2021, JrnPath [1] | |
| Basophil-Mast_1_DJ | 75 | 43 | Joseph, 2021, JrnPath | |
| Basophil-Mast_2_DJ | 75 | 42 | Joseph, 2021, JrnPath | |
| CD4+Memory-Effector_T_DJ | 75 | 45 | Joseph, 2021, JrnPath | |
| CD4+Naive_T_DJ | 75 | 37 | Joseph, 2021, JrnPath | CD4+ in Azimuth |
| CD8+Memory-Effector_T_DJ | 75 | 26 | Joseph, 2021, JrnPath | |
| CD8+Naive_T_DJ | 75 | 28 | Joseph, 2021, JrnPath | CD8+ in Azimuth |
| Classical_Monocyte_DJ | 75 | 32 | Joseph, 2021, JrnPath | |
| EREG+Dendritic_DJ | 75 | 41 | Joseph, 2021, JrnPath | |
| IGSF21+Dendritic_DJ | 75 | 41 | Joseph, 2021, JrnPath | |
| Interstitial_Fibroblasts_DJ | 75 | 38 | Joseph, 2021, JrnPath | |
| Myeloid_Dendritic_Type_1_DJ | 75 | 29 | Joseph, 2021, JrnPath | |
| Myeloid_Dendritic_Type_2_DJ | 75 | 35 | Joseph, 2021, JrnPath | |
| Natural_Killer_DJ | 75 | 21 | Joseph, 2021, JrnPath | |
| Natural_Killer_T_DJ | 75 | 22 | Joseph, 2021, JrnPath | |
| Nonclassical_Monocyte_DJ | 75 | 33 | Joseph, 2021, JrnPath | |
| Peri-epithelial_Fibroblasts_DJ | 75 | 38 | Joseph, 2021, JrnPath | |
| Pericytes_DJ | 75 | 45 | Joseph, 2021, JrnPath | |
| Plasma_DJ | 75 | 42 | Joseph, 2021, JrnPath | |
| Plasmacytoid_Dendritic_DJ | 75 | 24 | Joseph, 2021, JrnPath | |
| Proliferating_NK-T_DJ | 75 | 46 | Joseph, 2021, JrnPath | |
| Prostate_SM_DJ | 75 | 61 | Joseph, 2021, JrnPath | |
| TREM2+Dendritic_DJ | 75 | 61 | Joseph, 2021, JrnPath | |
| vSM_DJ | 75 | 38 | Joseph, 2021, JrnPath | |
| Basal_GH | 30 | 18 | Henry, 2018, CellRep [2] | |
| Club_GH | 69 | 56 | Henry, 2018, CellRep | |
| Endo_GH | 111 | 47 | Henry, 2018, CellRep | |
| Epithelia_GH | 166 | 56 | Henry, 2018, CellRep | |
| Fibroblast_GH | 124 | 61 | Henry, 2018, CellRep | |
| Hillock_GH | 54 | 30 | Henry, 2018, CellRep | |
| Leuk_GH | 135 | 65 | Henry, 2018, CellRep | |
| Luminal_GH | 71 | 63 | Henry, 2018, CellRep | |
| Neuro_GH | 560 | 55 | Henry, 2018, CellRep | |
| SmoothM_GH | 99 | 61 | Henry, 2018, CellRep | |
| Stress_GH | 13 | 13 | Henry, 2018, CellRep | |
| Stroma_GH | 905 | 96 | Henry, 2018, CellRep | |
| Stroma_RM | 150 | 42 | Tessem, 2016, PLOSOne [3] | |
| Citrate | 150 | 109 | Rye, 2022, iScience [4] | |
| GPS | 12 | *na* | Na, 2016, AsJrnAndr [5] | |
| Decipher | 19 | *na* | Na, 2016, AsJrnAndr | |
| CCP | 31 | *na* | Na, 2016, AsJrnAndr | |
| RA | 26 | 26 | This Study | |
| CEG | 12 | 12 | This Study | |

*Supplementary Table 3: Data sets used for analysis in public data. References below this Table.*

| ID | Dataset Abbreviation | Description | Data set source | Reference |
|---|---|---|---|---|
| 1 | Bertilsson | 156 prostate tissue samples (116 cancer and 40 normal) | Array Express: E-MTAB-1041 | [1] |
| 2 | Wang | 136 prostate tissue samples 65 cancer and 71 normal | GEO: GSE8218 | [2, 3] |
| 3 | Taylor | 160 prostate tissue samples (131 cancer and 29 normal) | GEO: GSE21034 | [4] |
| 4 | Sboner | 281 prostate cancer samples | GEO: GSE16560 | [5] |
| 5 | Erho | 545 prostate cancer samples | GEO: GSE46691 | [6] |
| 6 | TCGA-PRAD | 549 prostate tissue samples (497 cancer and 52 normal) | https://portal.gdc.cancer.gov/repository | https://www.cancer.gov/tcga |
| 7 | CMBR (Cambridge) | 186 prostate tissue samples (112 cancer and 74 normal) | GEO: GSE70768 | [7] |
| 8 | STCK (Stockholm) | 94 prostate cancer samples | GEO: GSE70769 | [7] |
| 9 | Mortensen | 50 prostate tissue samples (36 cancer and 14 normal) Laser dissected tissue | GEO: GSE46602 | [8] |
| 10 | Kuner | 98 prostate tissue samples (59 cancer and 39 normal) | GEO: GSE32571 | [9] |
| 11 | Loda | Laser Dissected tissue from 188 normal and cancer samples | GEO: GSE97284 | [10] |
| 12 | GTEx-Prostate | 245 prostate normal samples | https://gtexportal.org/home/datasets | https://gtexportal.org/home/ |
